# A novel inducible mtDNA mutator mouse model to study mitochondrial dysfunction with temporal and spatial control

**DOI:** 10.1101/2025.04.17.649454

**Authors:** Hannah Tobias-Wallingford, Sydney Bartman, Lauren Gaspar, Brian Gallagher, Carole Bassa, Emmanuel Sotirakis, Vincent Dubus, Christopher Hemme, Lars Olson, Giuseppe Coppotelli, Jaime M. Ross

**Author notes:** Correspondence should be addressed to: Jaime M. Ross Giuseppe Coppotelli. these authors contributed equally to this work (Sydney Bartman, Lauren Gaspar).

## Abstract

Mitochondrial dysfunction is a hallmark of aging and numerous age-related diseases. A wealth of studies supports the accumulation of mitochondrial DNA (mtDNA) mutations as a contributing factor to mitochondrial dysfunction in aging and disease. One of the best models to study the relationship between mtDNA mutations and mitochondrial dysfunction is the mtDNA mutator mouse, which expresses a proofreading-deficient version of mtDNA polymerase-γ (PolgA). Despite its groundbreaking contributions to mitochondrial biology and aging research, this model is limited by the whole-body accumulation of mtDNA mutations, which prevents the investigation of tissue-specific differences in mitochondrial dysfunction. To overcome this limitation, we developed a novel inducible knock-in mtDNA mutator mouse model that allows spatial and temporal control of mtDNA mutations, enabling the precise study of mitochondrial dysfunction in a tissue- and time-specific manner. Here, we report the generation and validation of this novel model through whole-body induction via Cre recombinase. Our data demonstrate that, upon induction, this model recapitulates the phenotype of the original mtDNA mutator mouse manifesting the same behavioral and biochemical alterations. This work establishes the functionality of our model and highlights its value as a powerful tool for studying the impact of mtDNA mutations with enhanced specificity and control.

## Introduction

The last several decades have seen significant improvements in healthcare, technology, and medicine, which has also brought a dramatic increase in life expectancy^1^. This change has resulted in shifting population dynamics, significantly increasing the percentage of individuals over the age of 65^2,3,4^. As a result, there has been a growing need for research to better understand aging and age-related diseases. While not fully understood, aging is described as the loss of biological processes and breakdown of organs, organelles, and macromolecules resulting in increased vulnerability to disease, and ultimately death^2,5,6^. To better understand and define aging, several key hallmarks were proposed and are now widely accepted, including genomic instability, telomere attrition, epigenetic alterations, loss of proteostasis, deregulation of nutrient sensing, mitochondrial dysfunction, cellular senescence, stem cell exhaustion, and alteration of intercellular communication^5,7,8^. Mitochondrial dysfunction, in particular, has repeatedly been identified as a major player in aging and many age-related diseases including mitochondrial diseases, cancers, and metabolic disorders^2^. Additionally, it has been implicated in neurodegenerative diseases, such as Alzheimer’s’, Parkinson’s’, and Huntington’s disease^9,10,11^. This inspired Denham Harman in 1972 to refine his 1956 Free Radical Theory of Aging, leading to the development of the Mitochondrial Theory of Aging^12,13^. The theory states that aging is due to slow accumulation of mitochondrial DNA (mtDNA) mutations over time, driven by reactive oxygen species (ROS), which disrupt bioenergetics and cellular homeostasis, leading to cellular dysfunction and cell death^12–14^.

Mitochondria are essential as the main energy producing organelle and play critical roles in calcium buffering, heme production, apoptosis, transcription, and several biosynthetic pathways vital for cell homeostasis and growth^3,15^. Mitochondria are the only organelles that possess their own DNA known as mitochondrial DNA (mtDNA), a circular molecule that encodes 13 essential components of the electron transport chain, including subunits of Complexes I, III, IV, and V. These complexes play a crucial role in the oxidative phosphorylation (OXPHOS) pathway, enabling ATP production within mitochondria^16^. mtDNA mutations can result in deficient OXPHOS activity and thus decreased ATP production^17^. Along with this, key signs of mitochondrial dysfunction include decreased mitochondrial membrane potential, swollen mitochondria, damaged cristae, increased oxidative stress, increased mtDNA mutations, and decreased mtDNA copy number^5,18^.

The mtDNA mutator mouse was one of the first models developed that allowed for studying the relationship between mtDNA mutations and aging. The model was first developed by the Nils- Gorän Larsson Laboratory at Karolinska Institutet in Sweden in 2004^19^, and independently by the Prolla Laboratory in the US in 2005^20^. The mtDNA mutator mouse is a knock-in homozygous model with a proofreading-deficient version (D257A) of the nuclear-encoded mtDNA polymerase-𝛾 (PolgA)^19,21^, which causes impaired proofreading ability thus resulting in an approximate linear accumulation of mtDNA mutations and deletions beginning during gestation. These mice exhibit a premature aging phenotype with features such as decreased lifespan, weight loss, decreased body size, sarcopenia, loss of subcutaneous fat, osteoporosis, kyphosis, alopecia, graying of the hair, decreased fertility, hearing loss, etc.^19,20,22,23^.

Since its development, the mtDNA mutator mouse model has provided some key insights into the role that mtDNA mutations and mitochondrial dysfunction have on health and aging. The model has been used by scientists interested in understanding the effect of mitochondrial dysfunction induced by mtDNA mutations on metabolism, insulin resistance, bone metabolism, intestine homeostasis, muscle waste, as well as Parkinson’s disease and related dementias^21,24–29^. Using this model, we have explored the role of germline mtDNA mutations and their impact on the aging process, brain development^22^, elevated lactate levels as a predictor of aging^30^, and the restoration of a dysregulated proteomic landscape through voluntary exercise^23^.

Despite the many seminal findings, the mtDNA mutator mouse model has several limitations that restrict its applicability. Since every cell in the body of this model experiences some level of mitochondrial dysfunction, all systems, including the visual and locomotor systems, are affected. As a result, studying cognition and behavior in this model is challenging, and distinguishing tissue and cellular differences in mitochondrial dysfunction becomes difficult. This makes it nearly impossible to fully understand the impact of mtDNA mutations on complex processes such as cognition, neural development, motor systems, and, more specifically, immune and inflammatory responses. To overcome these limitations, we developed an inducible model with temporal and spatial control of mitochondrial mutations. The aim of this report is to characterize this novel model and ensure recapitulation of the behavioral and cellular changes observed in the original mtDNA mutator mouse model, as well as to confirm the temporal and spatial control of mtDNA mutations.

## Results

### Generation of the inducible mtDNA mutator mouse model

The novel inducible mtDNA mutator mouse (i-PolgA^Mut/Mut^) enables inducible expression of the PolgA D257A mutation. To generate the i-PolgA^Mut/Mut^ model, the entire endogenous coding sequence (CDS) from exon 3 onwards was knocked-in along with the endogenous 3’UTR containing its polyadenylation signal followed by an exogenous polyadenylation signal derived from the human growth hormone (hGH polyA). This entire insertion is flanked by loxP sites. The same point mutation in exon 3 (G<u>A</u>C>G<u>C</u>C) used to generate the original mutator mouse^19^, which results in the D257A amino acid substitution in the protein, was introduced into exon 3 after the floxed cassette. This design ensures the expression of the wild-type PolgA allele until the CDS region is excised through Cre-mediated recombination (Fig. 1A and B).

**Figure 1.**
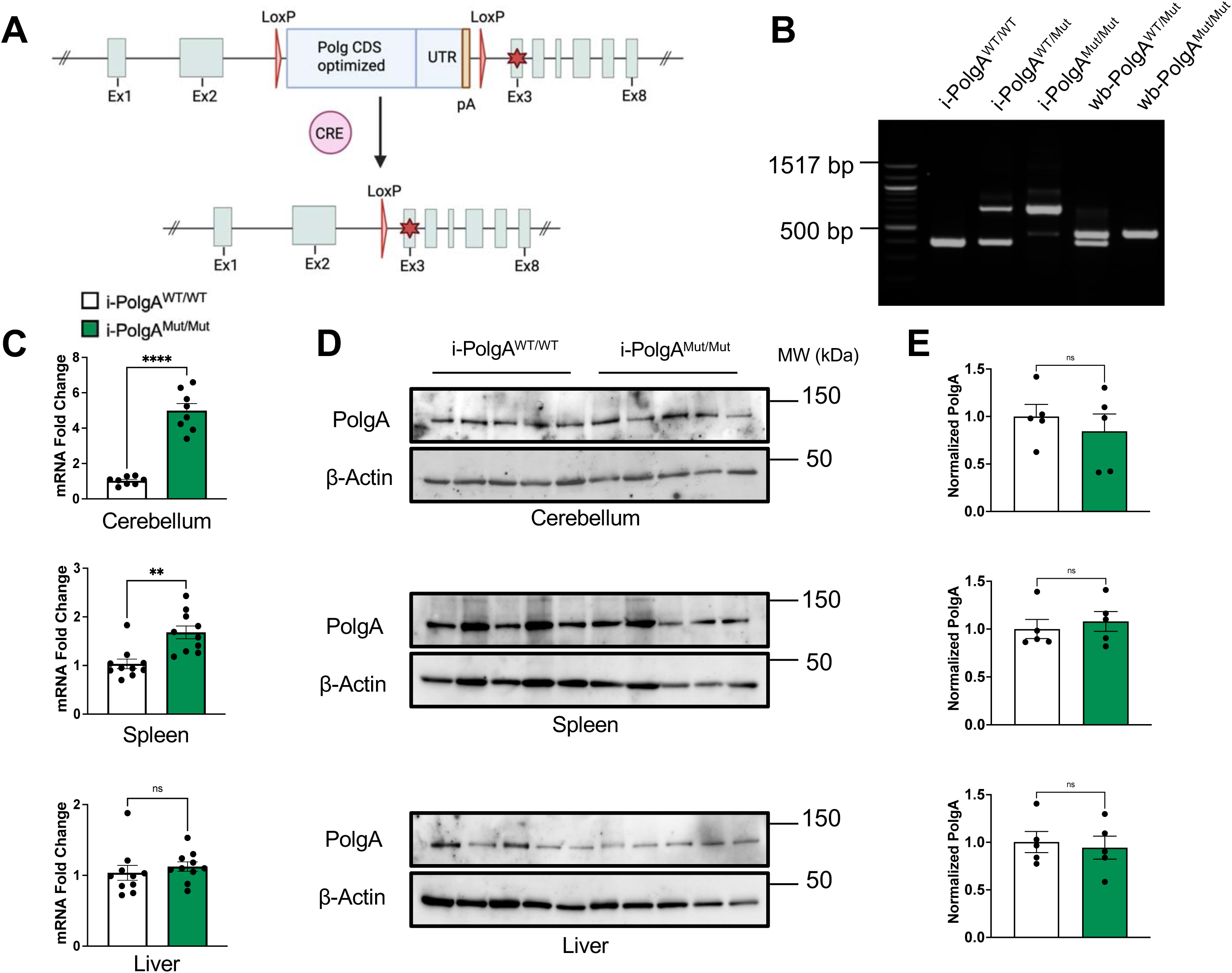
Generation and confirmation of the novel inducible mtDNA mutator mouse and PolgA expression. **(A)** Schematic of the Cre-loxP recombination strategy utilized to generate the novel PolgA inducible mutator mouse model (i-PolgA^Mut/Mut^). The PolgA CDS contains the entire CDS from exon 3 onwards, following this there is the endogenous untranslated region (UTR) and a hGH poly-adenylation (pA) signal (yellow). The mutation (red star) is within exon 3 and is only expressed following excision at the LoxP sites by CRE recombination. **(B)** Agarose gel electrophoresis of amplified PCR DNA products to visualize the novel PolgA inducible variants: i-PolgA^WT/WT^, i-PolgA^WT/Mut^, i-PolgA^Mut/Mut^, and the induced whole-body heterozygous wb- PolgA^WT/Mut^ and homozygous wb-PolgA^Mut/Mut^ mutator variants. **(C)** Real time PCR assessing PolgA mRNA expression in i-PolgA^WT/WT^ (male, N=10) and i-PolgA^Mut/Mut^ (male, N=10) animals across cerebellum, spleen, and liver tissue. **(D, E)** Western blot analysis to assess protein levels of PolgA across tissues from i-PolgA^WT/WT^ (male, N=5) and i-PolgA^Mut/Mut^ (male, N=5) animals. Significances were determined by unpaired t-test **(C and E)**; denoted ** *p* < 0.01, **** *p* < 0.0001.

### Characterization of the inducible mtDNA mutator mouse, i-PolgA^Mut/Mut^

In order to determine that the insertion of the CDS cassette in the PolgA locus was not affecting this novel model, we verified that there were no unexpected differences in the uninduced i- PolgA^Mut/Mut^ animals compared to wild-type control littermates (i-PolgA^WT/WT^). We started by examining PolgA expression and protein levels in multiple tissues from i-PolgA^Mut/Mut^ mice. Although qPCR analysis found tissue specific variation in mRNA expression of PolgA (Fig. 1C), Western blot analysis showed similar protein levels of PolgA in liver, spleen and cerebellum from i-PolgA^WT/WT^ and i-PolgA^Mut/Mut^ animals (Fig. 1D and E), thus suggesting that the cassette has no major effect on the PolgA protein levels in the i-PolgA^Mut/Mut^ model.

Using the frailty index aging scoring system to assess overall phenotypes, we found no differences in the i-PolgA^Mut/Mut^ mice as compared to controls (Fig. 2A, B). We also found no differences in body weights of i-PolgA^Mut/Mut^ mice over the course of forty weeks (Fig. 2C). Both heterozygous i-PolgA^WT/Mut^ and homozygous i-PolgA^Mut/Mut^ mice were fertile with no deviations in fertility and fecundity found (Fig. 2D left). Moreover, intercrosses of i-PolgA^WT/Mut^ mice produced offspring born in a Mendelian fashion (Fig. 2D right). Gross examination of peripheral organs revealed no macroscopic differences, and no significant differences were observed in the percentage of body weight for the heart, liver, and spleen (Fig. 2E).

**Figure 2.**
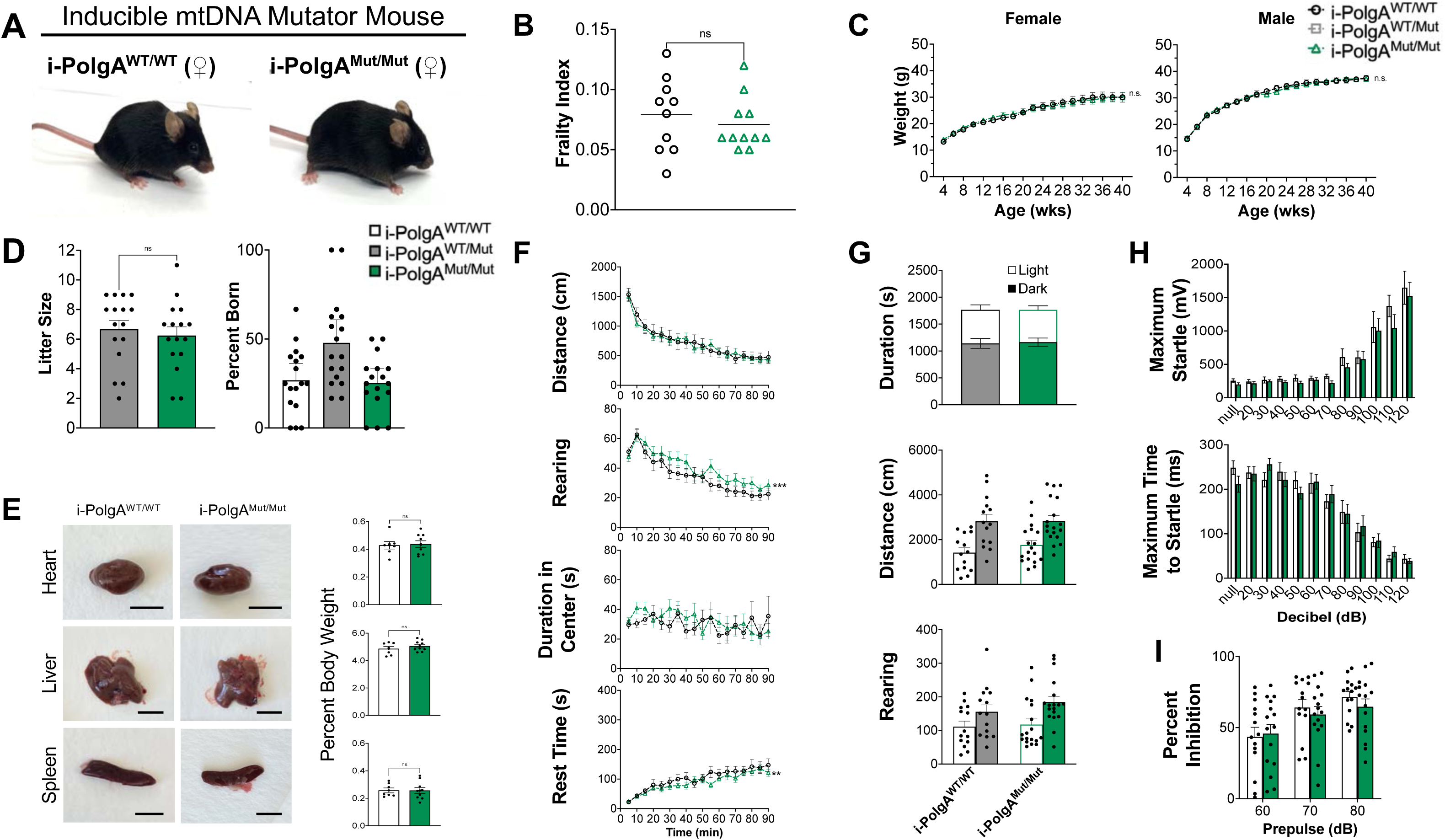
Characterization of the inducible mutator model, i-PolgA^Mut/Mut^. **(A)** Representative images of an uninduced i-PolgA^Mut/Mut^ mouse at 36 weeks of age showing no phenotype, as compared to an age-matched i-PolgA^WT/WT^ control mouse (females shown). **(B)** Scores obtained from frailty index (FI) testing of 36 week-old i-PolgA^WT/WT^ (N=10) and i-PolgA^Mut/Mut^ (N=10) mice. **(C)** Body weight measurements for i-PolgA^WT/WT^ (black) and i-PolgA^Mut/Mut^ (green) mice beginning at 4 to 40 weeks of age, taken every 2 weeks. Graphs shown are female (N=10 i- PolgA^WT/WT^ and N=10 i-PolgA^Mut/Mut^) and male (N=10 i-PolgA^WT/WT^ and N=11 i-PolgA^Mut/Mut^) animals. **(D)** Graph on the left shows litter sizes from intercrosses of i-PolgA^WT/Mut^ mice (gray) and intercrosses of i-PolgA^Mut/Mut^ mice (green) while on the right shows the percentage of i- PolgA^WT/WT^, i-PolgA^WT/Mut^, and i-PolgA^Mut/Mut^ born from intercrosses of i-PolgA^WT/Mut^ mice. **(E)** Representative images of heart, liver, and spleen from 40 week-old female i-PolgA^WT/WT^ and i- PolgA^Mut/Mut^ mice, scale bar=1 cm. Percent of body weight of heart, liver, and spleen, respectively, in i-PolgA^WT/WT^ and i-PolgA^Mut/Mut^ mice. **(F)** Spontaneous locomotor and exploratory activity of 36 week-old i-PolgA^WT/WT^ (N=14) and i-PolgA^Mut/Mut^ (N=18) mice in the open-field assay. **(G)** Exploratory and anxiety-like behavior in the light-dark preference assay in 36 week-old i- PolgA^WT/WT^ (N=14) and i-PolgA^Mut/Mut^ (N=18) mice. **(H)** Maximum startle (mV) as well as maximum time to startle (ms) were measured in 36 week-old i-PolgA^WT/WT^ (N=14) and i- PolgA^Mut/Mut^ mice (N=18). **(I)** Percent pre-pulse inhibition in the startle reflex assay in 36 week-old i-PolgA^WT/WT^ (N=14) and i-PolgA^Mut/Mut^ mice (N=18). Significances were determined by two-way ANOVA with post-hoc analysis (**A, B, C, D)**; denoted ** *p* < 0.01, *** *p* < 0.001.

Behavioral testing was also performed on 9 months old i-PolgA^Mut/Mut^ mice and i-PolgA^WT/WT^ littermate controls. Spontaneous locomotor movements and exploratory behavior were monitored in environmentally controlled chambers for 90 minutes while distance traveled, rearing, duration in center/periphery, and rest time were measured. No differences in distance traveled and duration in center were found in i-PolgA^Mut/Mut^ animals compared to i-PolgA^WT/WT^ controls (Fig. 2F, Fig. S1A). No differences in total rearing activity and total rest time in i-PolgA^Mut/Mut^ animals were found (Fig. S1A), but an increase in rearing activity and a decrease in rest time were observed when comparing 5-minute intervals over the 90 minutes (Fig. 2F), due to some variability in sex differences. (Fig. S1B). Specifically, i-PolgA^Mut/Mut^ females showed an increase in rearing activity and duration spent in the center, whereas i-PolgA^Mut/Mut^ males showed a slight decrease in rest time (Fig. S1B).

Anxiety-like behavior was assessed using the light-dark preference test, in which the open-field chamber was divided into light and dark zones, and the time spent in each zone was monitored to determine preference by measuring distance traveled, rearing activity, and resting time in each location. No differences were found between i-PolgA^Mut/Mut^ mice and i-PolgA^WT/WT^ controls, and both groups performed as expected by spending more time and moving more in the dark compartment as compared to the light one (Fig. 2G, S2A and B). Both i-PolgA^Mut/Mut^ mice and i- PolgA^WT/WT^ controls had expected startle reflex and no changes in maximum startle reflex and maximum time to startle parameters were found between groups (Fig. 2H). Lastly, both mouse types showed expected percent inhibition during the pre-pulse inhibition assay, and no differences were found between the i-PolgA^Mut/Mut^ and i-PolgA^WT/WT^ animals (Fig. 2I).

Lastly, we confirmed no changes in mitochondrial function in the i-PolgA^Mut/Mut^ mice compared to the i-PolgA^WT/WT^ control. This was determined by assessing Complex IV activity using cytochrome c oxidase (COX) staining and Complex I activity in liver tissue (Fig. 7B and C).

Additionally, there were no differences in mtDNA copy number in skeletal muscle or mtDNA mutation load and mutation type in liver tissue between the i-PolgA^Mut/Mut^ and i-PolgA^WT/WT^ mice (Fig. 7D-H). Finally, no differences in lifespan were observed between i-PolgA^Mut/Mut^ and i- PolgA^WT/WT^ mice up to 90 weeks of age (Fig. 7I).

### The whole-body induced PolgA mutator mouse recapitulates the original mtDNA mutator model

To validate the new i-PolgA^Mut/Mut^ model as an inducible mtDNA mutator mouse, we tested whether whole-body induction via Cre recombination (wb-PolgA^Mut/Mut^) (Fig. 3E) would recapitulate the phenotype of the original mtDNA mutator mouse (PolgA^Mut/Mut^) model (Fig. 3A). Aging phenotypes were assessed utilizing frailty index aging scoring and monitoring body weights in homozygote WT, heterozygote, and homozygote Mut animals in both the novel wb-PolgA and original PolgA models (Fig. 3B, C, F, G). Previous work has shown that the original mtDNA mutator mouse exhibits multiple signs of premature aging with decreased body size, kyphosis, alopecia, osteoporosis, and enlarged heart, spleen, and liver^19^, and we found that the novel induced wb-PolgA mutator mouse (Fig. 3E) shows visible signs of premature aging similar to the original mtDNA mutator mouse (Fig. 3A), such as decreased body size, graying hair, alopecia, and kyphosis. Body weight trends from 4 to 40 weeks of age in wild-type, heterozygous, and mutator animals in both models were also similar and in line with the original mtDNA mutator model, with PolgA^Mut/Mut^ mice (Fig. 3C) and wb-PolgA^Mut/Mut^ mice (Fig. 3G) slowing in weight gain around 15-20 weeks of age and losing weight from about 24 weeks onwards. Lastly, wild-type, heterozygous, and mutator animals of both models were blindly rated using a 31-point frailty index scale to further confirm these trends, with a higher frailty index score indicating a frailer phenotype. Both the original (Fig. 3B) and the novel induced (Fig. 3F) mutator models had significantly higher frailty index scores at 36 weeks of age than their age-matched wild-type and heterozygous counterparts. The heart, liver, and spleen were significantly enlarged in the mutator animals of both models, accounting for a greater percentage of body weight in the novel induced wb-PolgA mutator mouse and the original mtDNA mutator mouse, compared to wild-type controls (Fig. 3D and H).

**Figure 3.**
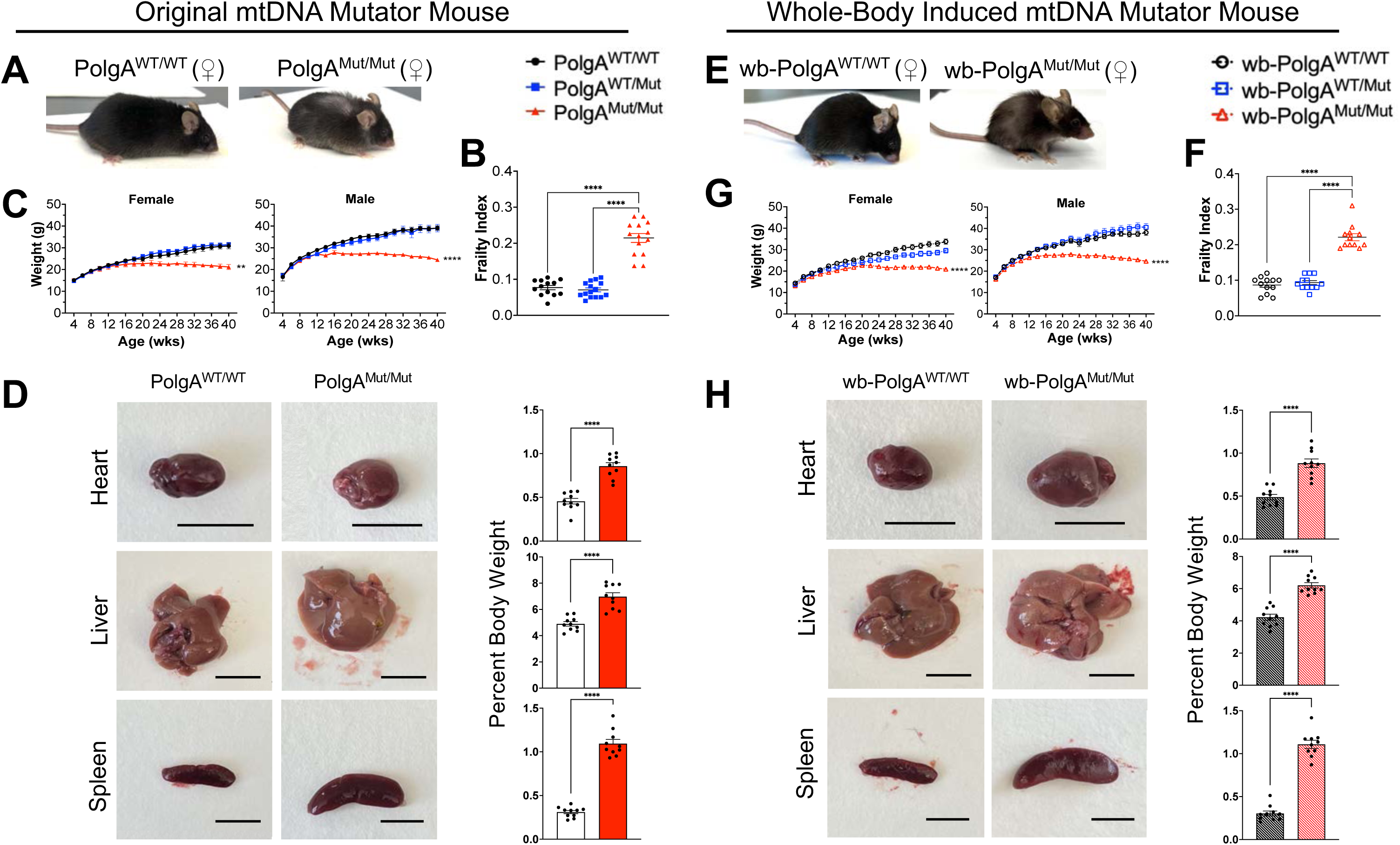
Characterization of the novel inducible mtDNA mutator mouse compared to the original mtDNA mutator model. **(A)** Representative images of a PolgA^Mut/Mut^ mouse at 40 weeks of age showing progeroid phenotypes of decreased body size, alopecia, graying of the hair, kyphosis, etc., as compared to an age-matched PolgA^WT/WT^ littermate (females shown). **(B)** Frailty index (FI) aging scores of 36 week-old PolgA^WT/WT^ (N=13), PolgA^WT/Mut^ (N=16), and PolgA^Mut/Mut^ (N=13) animals. **(C)** Body weight measurements of 4 to 40 week-old PolgA^WT/WT^, PolgA^WT/Mut^, and PolgA^Mut/Mut^ female (N= 10 per group) and male (N=10 per group) mice, taken every 2 weeks. **(D)** Representative images of heart, liver, and spleen from 40 week-old PolgA^WT/WT^ and PolgA^Mut/Mut^ mice, scale bar = 1 cm. Percent of body weight of heart, liver, and spleen, respectively (N=10 per group). **(E)** Representative image of a wb-PolgA^Mut/Mut^ mouse at 40 weeks of age with an age-matched wb-PolgA^WT/WT^ littermate (females shown), with the wb-PolgA^Mut/Mut^ model showing similar progeroid characteristics to the PolgA^Mut/Mut^ model described previously, including decreased body size, alopecia, graying of the hair, kyphosis, etc. **(F)** FI scores of 36 week-old wb-PolgA^WT/WT^ (N=12), wb-PolgA^WT/Mut^ (N=12), and wb-PolgA^Mut/Mut^ (N=12) animals. **(G)** Body weight curves of 4 to 40 week-old wb-PolgA^WT/WT^, wb-PolgA^WT/Mut^, and wb- PolgA^Mut/Mut^ females (N=12, 12, 10, respectively) and males (N=10, 10, 11, respectively), taken every 2 weeks. **(H)** Images of heart, liver, and spleen from 40 week-old wb-PolgA^WT/WT^ and wb- PolgA^Mut/Mut^ mice, scale bar = 1 cm. Percent of body weight of heart, liver, and spleen, respectively (N=10 per group). Significances were determined by RM two-way ANOVA **(C, G)**, two-way ANOVA **(B, F)**, or one-way ANOVA **(D, H)**, all tests with post-hoc analyses; denoted ** *p* < 0.01, **** *p* < 0.0001.

### Behavioral characterization of the whole-body induced mtDNA mutator mouse are comparable to the original mtDNA mutator mouse model

Further characterization of the novel induced wb-PolgA mutator mouse and comparison to the original mtDNA mutator mouse was done through behavioral assays including open-field (Fig. 4 and S3-S7), light/dark preference (Fig. 5 and S8-12), startle reflex (Fig. 6A), and pre-pulse inhibition (Fig. 6B). Open-field and light/dark preference were performed at 3, 6, and 9 months in both models. Changes in exploratory behavior and spontaneous locomotion were observed at each time-point, with the most drastic changes at 9 months. Both PolgA^Mut/Mut^ and wb-PolgA^Mut/Mut^ mice showed decreases in heat map signatures, distance traveled, and rearing activity as well as increases in rest time, as compared to controls (Fig. 4 and S3). No striking sex differences were found in the novel and original mutator models (Fig. S4-7). Similar changes were observed in the light/dark preference test, confirming the open-field findings (Fig. 5, S8) with no marked sex differences observed (Fig. S9-12). The startle reflex response was tested at 6 and 9 months in both the original and novel models and both PolgA^Mut/Mut^ and wb-PolgA^Mut/Mut^ mice showed a decreased startling to lower, but not higher, decibels at 9 months of age (Fig. 6A), which could be due to hearing loss. At 9 months, the pre-pulse inhibition assay was performed in both models and no significant differences were observed in the PolgA and the wb-PolgA models when comparing mutator with wild-type animals, while a minor decrease was found in wb-PolgA heterozygous animals (Fig. 6B).

**Figure 4.**
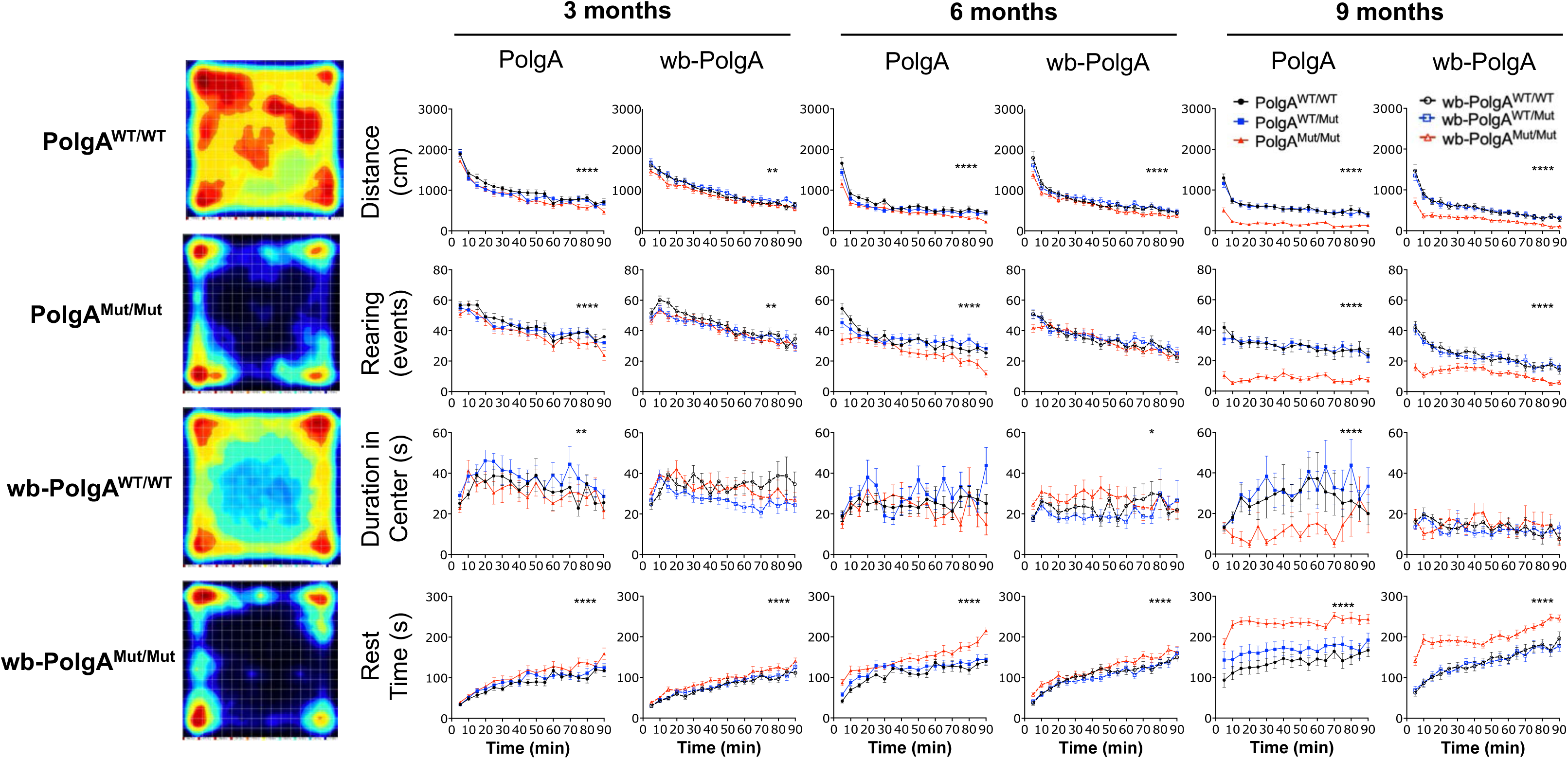
Open-field behavioral assay in novel wb-PolgA mice as compared to orginal PolgA model. Spontaneous locomotion and exploratory activity in the original PolgA^WT/WT^ (N=20), PolgA^WT/Mut^ (N=20), and PolgA^Mut/Mut^ (N=20) mice, as compared to the novel wb-PolgA^WT/WT^ (N=22), wb-PolgA^WT/Mut^ (N=23), wb-PolgA^Mut/Mut^ (N=24) animals at 3, 6, and 9 months of age. Both PolgA^Mut/Mut^ and wb-PolgA^Mut/Mut^ mice exhibited altered behavior including decreased distance traveled, decreased rearing, and increased rest time, all parameters which worsened with increased age (right). Representative heat map (left) illustrating the time spent by the mouse in different locations of the testing arena during the assay. Significances were determined by two-way ANOVA with post-hoc analysis, and denoted by * *p* < 0.05, ** *p* < 0.01, *** *p* < 0.001, **** *p* < 0.0001.

**Figure 5.**
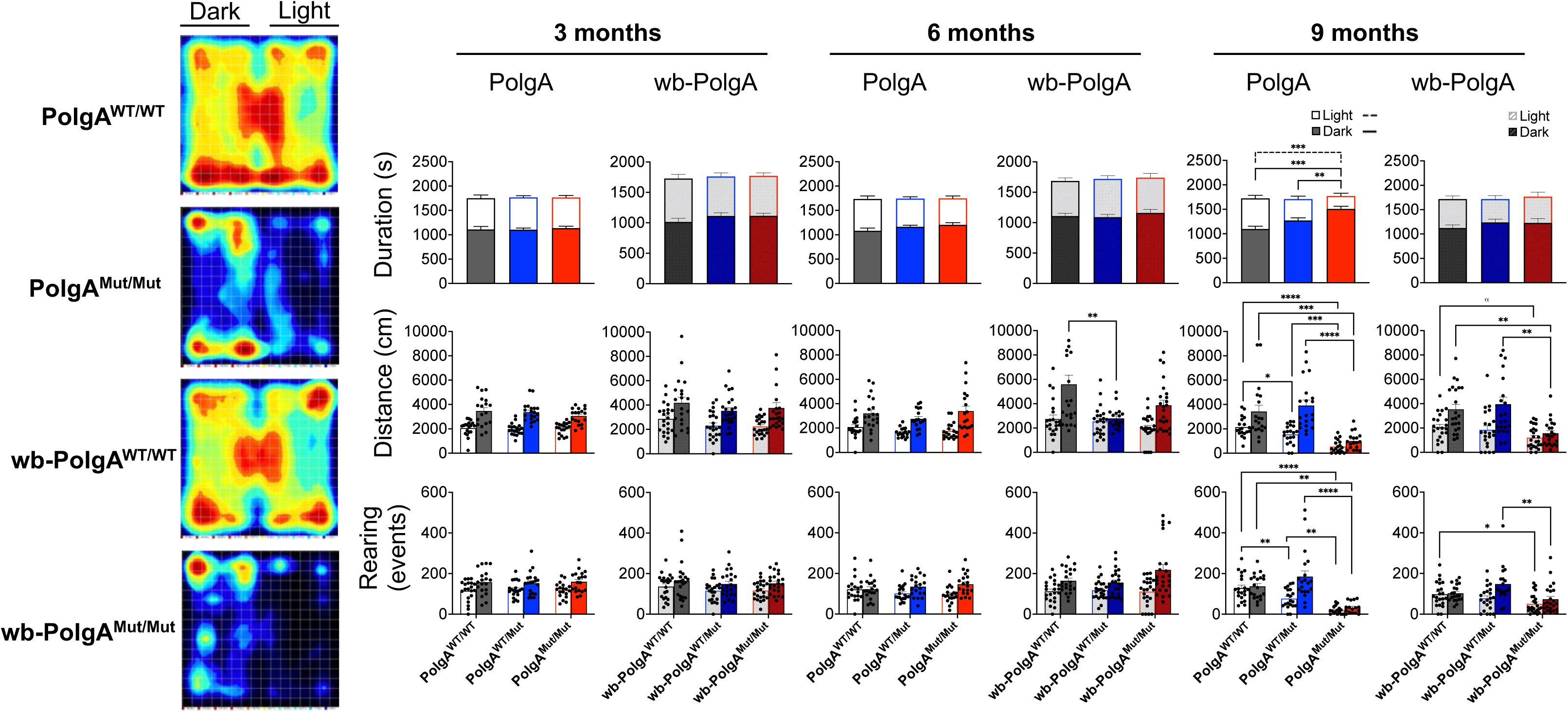
Light-dark behavioral assay in PolgA and wb-PolgA mice. Exploratory and anxiety-like behaviors were examined using the light-dark preference assay in the novel wb-PolgA^WT/WT^ (N=22), wb-PolgA^WT/Mut^ (N=23), and wb-PolgA^Mut/Mut^ (N=24) mice as compared to the original PolgA^WT/WT^ (N=20), in PolgA^WT/Mut^ (N=20), PolgA^Mut/Mut^ (N=20) model at 3, 6, and 9 months of age. Both PolgA^Mut/Mut^ and wb-PolgA^Mut/Mut^ animals exhibited altered behavior including decreased rearing and distance traveled (right). Representative heat map (left) illustrating the time spent by the mouse in different locations of the testing arena during the assay. Significances were determined by one-way ANOVA with post-hoc analysis; denoted as * *p* < 0.05, ** *p* < 0.01, *** *p* < 0.001, **** *p* < 0.0001, with α *p* < 0.10 as trending.

**Figure 6.**
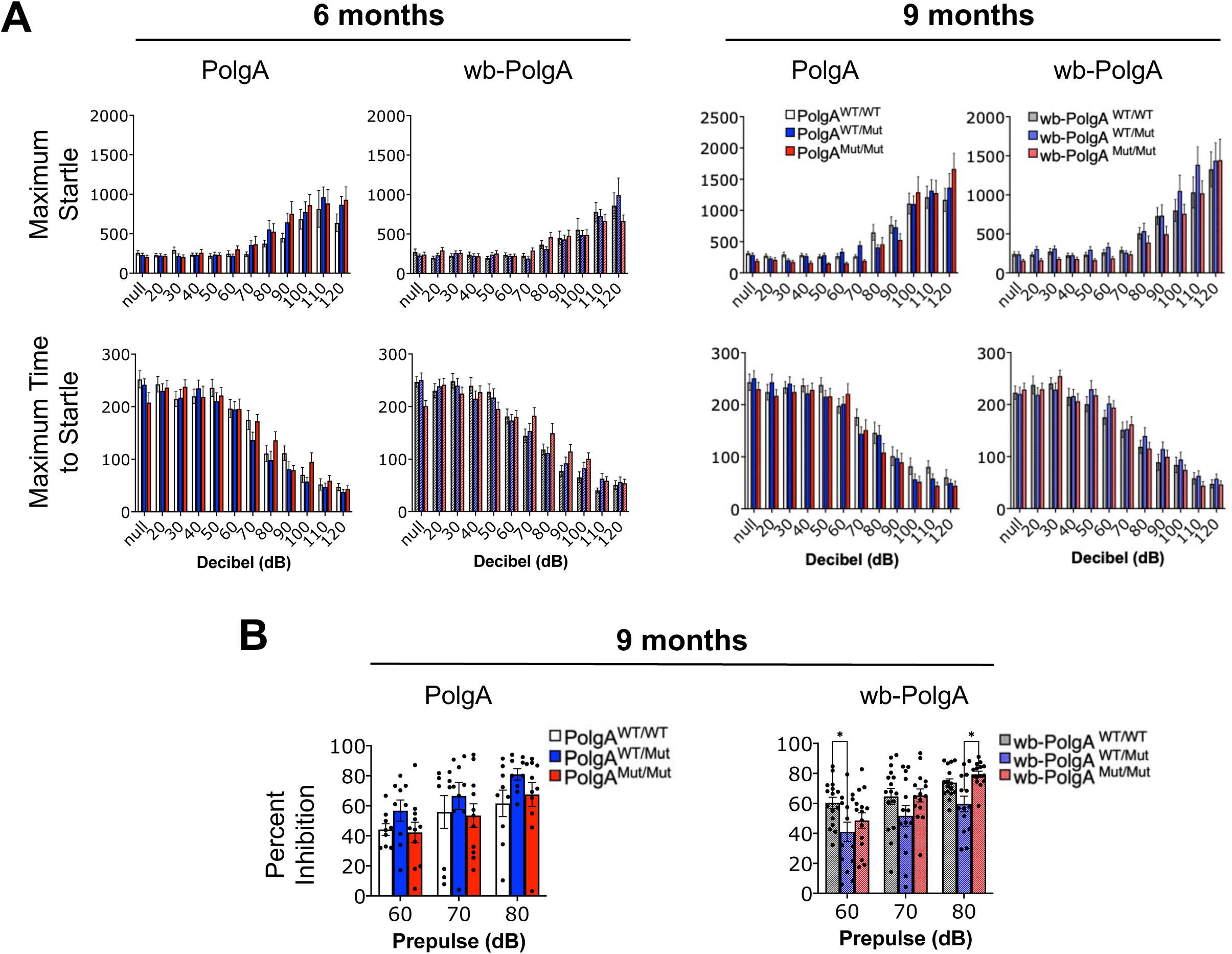
Startle reflex and pre-pulse inhibition behavioral assay in PolgA and wb-PolgA mice. **(A)** Startle reflex behaviors in 6 and 9 month-old novel wb-PolgA^WT/WT^ (N=22), wb- PolgA^WT/Mut^ (N=23), and wb-PolgA^Mut/Mut^ (N=24) mice, as compared to the original PolgA^WT/WT^ (N= 20), PolgA^WT/Mut^ (N=20), and PolgA^Mut/Mut^ (N=19) model. **(B)** Pre-pulse inhibition (PPI) in 9 month-old novel wb-PolgA^WT/WT^ (N=17), wb-PolgA^WT/Mut^ (N=16), and wb-PolgA^Mut/Mut^ (N=17) mice, as compared to the original PolgA^WT/WT^ (N=10), PolgA^WT/Mut^ (N=10), and PolgA^Mut/Mut^ (N=13) model. Significances were determined by two-way ANOVA with post-hoc analysis and denoted as * *p* < 0.05, ** *p* < 0.01.

### Mitochondrial structural and functional changes in both mutator models

Transmission electron micrographs of gastrocnemius from 9 months old wb-PolgA^Mut/Mut^ mice revealed degeneration of muscle fiber architecture, along with altered mitochondrial morphology, cristae structure, and orientation compared to age-matched controls (Fig. 7A). These findings are consistent with observations in the original mtDNA mutator mouse^19^. To further assess mitochondrial function, respiratory enzyme function was evaluated using cytochrome c oxidase (COX) histochemistry. A reduction in COX activity was observed in 9 months old PolgA^Mut/Mut^ and wb-PolgA^Mut/Mut^ liver samples, indicated by fainter staining compared to controls (Fig. 7B). Additionally, liver extracts from both PolgA^Mut/Mut^ and wb-PolgA^Mut/Mut^ mice showed reduced Complex I activity compared to wild-type controls (Fig. 7C). Furthermore, both PolgA^Mut/Mut^ and wb-PolgA^Mut/Mut^ mice at 9 months exhibited a decrease in mtDNA copy number in gastrocnemius relative to age-matched controls (Fig. 7D).

**Figure 7.**
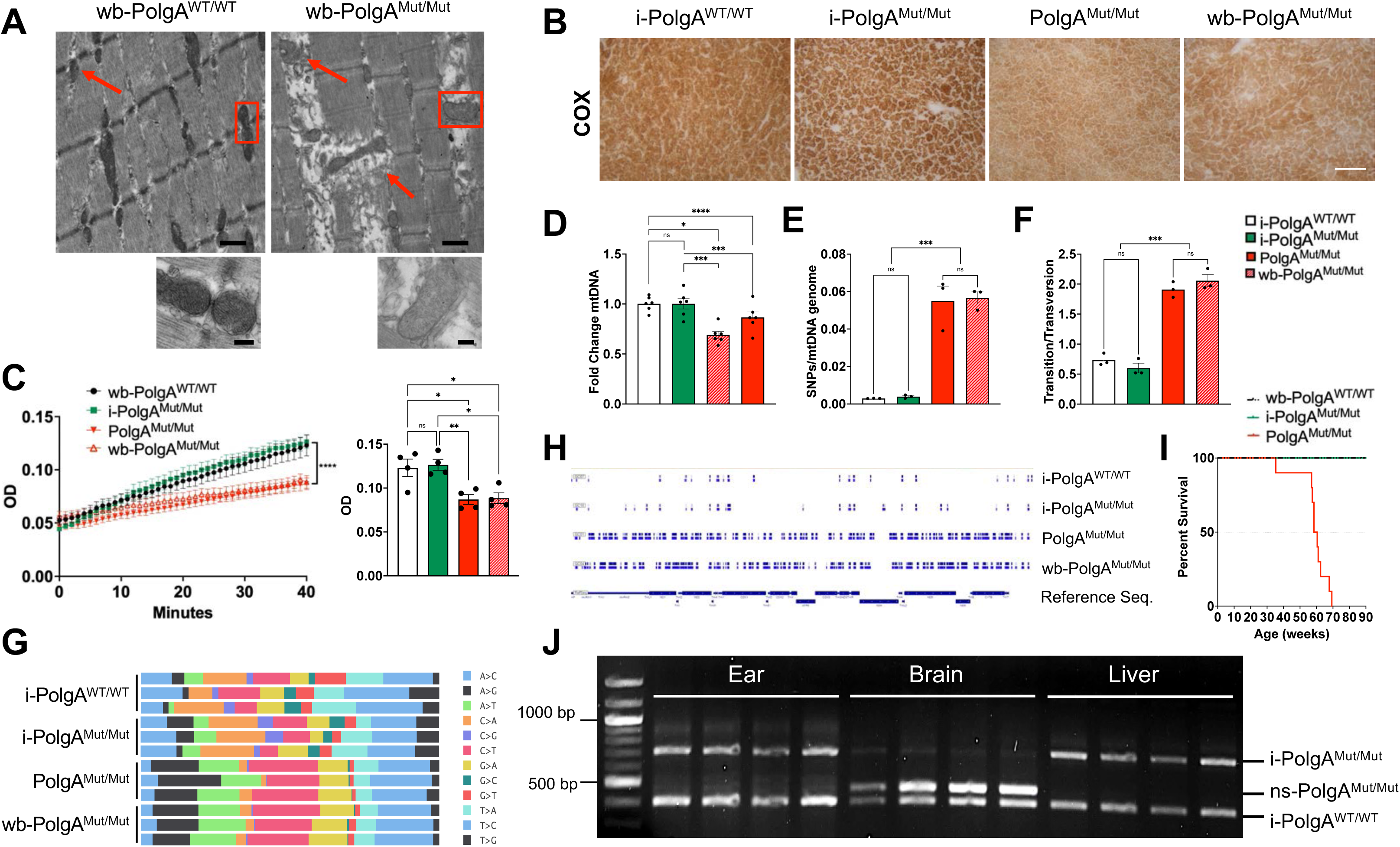
Mitochondrial structural and functional changes in wb-PolgA^Mut/Mut^ animals. **(A)** Representative images of transmission electron micrographs of gastrocnemius muscle from 40 week-old wb-PolgA^WT/WT^ and wb-PolgA^Mut/Mut^ animals, with mitochondria indicated by red arrows and enlargement in red boxes. Changes in mitochondrial morphology and orientation were observed in both the original and novel mutator models (male, N=3 observed per group). **(B)** Representative images showing decreased cytochrome *c* oxidase (COX) enzyme histochemistry in liver sections from 40-week-old PolgA^Mut/Mut^ and wb-PolgA^Mut/Mut^ mice, as compared to male i- PolgA^WT/WT^ and i-PolgA^Mut/Mut^ mice (male, N=3 per group). **(C)** Decreased mitochondrial Complex I activity in liver from 40-week-old PolgA^Mut/Mut^ and wb-PolgA^Mut/Mut^ mice, as compared to i-PolgA^WT/WT^ and i-PolgA^Mut/Mut^ mice (female, N=4 per group). **(D)** Decreased mitochondrial copy number assessed by qPCR as well as **(E)** increased mtDNA mutation load and **(F)** increased transition/transversion ratio assessed by next-generation sequencing (NGS) were found in PolgA^Mut/Mut^ and wb-PolgA^Mut/Mut^ mice, as compared to i-PolgA^WT/WT^ and i-PolgA^Mut/Mut^ animals (male, 40 weeks old, N= 3 per group). **(G)** Visualization of spread of mtDNA mutation type in PolgA^Mut/Mut^ and wb-PolgA^Mut/Mut^, as compared to i-PolgA^WT/WT^ and i-PolgA^Mut/Mut^ mice (male, 40 weeks old, N=3 per group). **(H)** Visual representation of the total mtDNA mutation load in PolgA^Mut/Mut^ and wb-PolgA^Mut/Mut^, as compared to i-PolgA^WT/WT^ and i-PolgA^Mut/Mut^ mice, with blue lines indicating changes from the reference sequence (bottom). **(I)** Kaplan-Meier graph showing decreased lifespan of wb-PolgA^Mut/Mut^ mice in comparison to wb-PolgA^WT/WT^ and i-PolgA^Mut/Mut^ mice (females, N=10 per group). **(J)** Agarose gel electrophoresis of PCR DNA products confirming brain specific induction of the i-PolgA^Mut/Mut^ following cross with Nestin-Cre mouse model, the ns-PolgA^Mut/Mut^ mouse (N=4). Significances were determined by two-way ANOVA and one-way ANOVA, both with post-hoc analyses **(C, D),** or unpaired two-tailed and one-tailed t-tests **(D)**; denoted as \**p* < .05, \*\**p* < .01, \*\*\**p* < .001, \*\*\*\**p* < .0001.

### Elevated mtDNA mutation load in both mutator models

To assess mtDNA mutation levels, we performed Next-Generation Sequencing (NGS) on mtDNA extracted from liver samples from both the novel and original mutator models, as well as appropriate controls, at 9 months of age. NGS analysis revealed elevated mtDNA mutation loads in both the original PolgA^Mut/Mut^ and wb-PolgA^Mut/Mut^ mice with similar mutation locations (Fig. 7E and H). Additionally, a similar increase in transition/transversion ratio was found in both the original and novel mutator models, as compared to controls (Fig. 7F). Analysis also revealed that the type of mtDNA mutation shifts were similar in both the novel and original mutator models (Fig. 7G).

### Shortened lifespan observed in both mutator models

We observed that wb-PolgA^Mut/Mut^ mice had an average lifespan of ∼55 weeks and a maximal lifespan of ∼65 weeks, significantly reduced compared to wb-PolgA^WT/WT^ and i-PolgA^Mut/Mut^ mice (Fig. 7I). This shorted lifespan is comparable to what was reported in the original mtDNA mutator mouse model (median lifespan of ∼48 weeks and maximal ∼61 weeks)^19^.

### Confirmation of tissue-specific induction of the i-PolgA^Mut/Mut^ mouse

Tissue specificity of the novel i-PolgA^Mut/Mut^ mouse was confirmed utilizing a Nestin-Cre model that expresses Cre recombinase under control of the Nestin promoter, which is expressed in the central and peripheral nervous systems^31,32^. Offspring obtained from crossing the Nestin-Cre and i-PolgA^Mut/Mut^ strains were genotyped, revealing tissue-specific recombination in the brain but not in liver tissue or ear biopsies from the nervous-system specific mtDNA mutator mouse (nsPolgA^Mut/Mut^) (Fig. 7J).

## Discussion

The mtDNA mutator mouse model has been a critical tool in assessing physiological impacts of mitochondrial dysfunction; however, its use has been limited to studying whole-body and life-long accumulation of mtDNA mutations. In this technical report, we introduce the first novel mouse model that has been developed to allow spatial and temporal control of mtDNA mutations. Given the rise in aging populations and strong association between mitochondrial dysfunction and many age-related diseases, we believe it is essential to have a model to address important questions regarding tissue and time-point specific mtDNA mutation effects. Thus, we sought to establish the credibility of this novel model, ensure recapitulation of key features of the original mtDNA mutator mouse, and provide a robust baseline behavioral and phenotypic characterization to guide future use.

Other “inducible” versions of the original mtDNA mutator mouse have been attempted in recent years^33,34^; however, the phenotypes in these new models were either unverified^33^ or did not recapitulate^34^ the original model. For example, in the model by Kubekina et al., a transgene carrying a mutated PolgA allele downstream of a floxed stop cassette was used to generate a transgenic mouse via pronuclear injection, allowing for the expression of mutated PolgA upon Cre recombination. In this model, the continuous expression of the endogenous PolgA gene most likely prevents any phenotype from manifesting, similar to what has been observed in the heterozygous PolgA mutator mouse^19,26^. The most recent model by Bond et al. utilizes Cre recombination to delete Exons 4 and 5, resulting in a partial deletion of the exonuclease domain. However, the stability and activity of the truncated PolgA in this model were never demonstrated. Additionally, the model did not show an increase in mtDNA mutation load in a muscle-specific context, and no attempts were made to assess whether whole-body deletion of the domain would replicate the original mutator mouse phenotype.

To characterize our novel model, we first wanted to ensure that the uninduced mouse, i- PolgA^Mut/Mut^, developed similarly to a wild-type animal and that the CDS cassette did not leak. Phenotype was evaluated using frailty index scoring, body weight was assessed from 4 until 40 weeks of age in both male and female mice, and organ weights with percentage of body weight was calculated, with no differences identified (Fig. 2A, B, C, E). Additionally, typical fertility and fecundity was observed in both heterozygous and homozygous crosses (Fig. 2D). i-PolgA^Mut/Mut^ animals were tested in a series of behavioral assays including open-field, light/dark preference test, startle reflex, and pre-pulse inhibition, and these assays showed no or limited differences in behavior in the uninduced mouse, compared to wild-type mice (Fig. 2F, G, H). Additionally, while PolgA mRNA varied in qPCR analyses (Fig. 1C), there were no differences in PolgA protein levels between i-PolgA^Mut/Mut^ and i-PolgA^WT/WT^ animals in western blot analyses (Fig. 1D and E). No differences in mitochondrial function between the uninduced and wild-type animals were found in COX histochemistry (Fig. 7B) or in Complex I activity (Fig. 7C). Additionally, there was no difference in mtDNA copy number, a biomarker of mitochondrial function, between the wild-type and uninduced animals (Fig. 7D). Uninduced and wild-type animals were observed until 90 weeks of age with no differences in lifespan found (Fig. 7I). Most importantly, there was no increase in mtDNA mutation load in the uninduced animals and the mtDNA mutations were similar to wild-type animals in frequency and type (Fig. 7E-H). Taken together, these findings suggest that the uninduced mouse, i-PolgA^Mut/Mut^ develops typically, does not show any differences from its wild-type littermate, and importantly appears to have no change in mitochondrial function or mitochondrial genome due to a possible leakage of the transgene expression.

Before embarking in any tissue- or time-specific analysis of mtDNA mutations, we first aimed to confirm that whole-body induction of the novel i-PolgA^Mut/Mut^ mouse would recapitulate the original mtDNA mutator mouse model both genotypically and phenotypically, given that both models have the same point mutation in Exon 3 (D257A) of the mtDNA proofreading polymerase-𝛾 that results in mtDNA mutations and deletions that accumulate from gestation. Past work has shown that the original mutator mouse has decreased body weight, several phenotypic traits of premature aging, and enlargement of the heart, liver, and spleen^19^. Both the original mtDNA mutator mouse, PolgA^Mut/Mut^, and the novel whole-body induced mutator mouse, wb-PolgA^Mut/Mut^, presented with similar increases in frailty as well as other premature aging phenotypes such as alopecia, kyphosis, graying of hair, decreases in body weight, enlargement of heart, liver, and spleen (Fig. 3D and H), as well as premature death (Fig. 7I). Using several behavioral assays, both models exhibited similar trends, with decreased distance traveled and rearing activity, as well as increased resting in both the open-field and light/dark preferences tests (Fig 5 and 6). Although there were no clear differences in startle reflex, mice harboring the mutated polymerase-𝛾 appeared to begin startling only at higher decibel levels (Fig. 7), possibly due to hearing loss, as previously reported^35^.

Alterations in mitochondrial function and morphology were observed in both the original and novel whole-body induced mutator models. TEM images of gastrocnemius showed altered mitochondria morphology, cristae, and orientation alongside degradation of the muscle fiber architecture in the induced mutator mouse (Fig. 7A), similar to what has been reported in the original model^19,23^. Mitochondrial function assessment revealed similar deficient respiratory activities of Complexes I and IV, as determined by COX enzyme histochemistry and activity assay (Fig. 7B and C). Additionally, qPCR analysis revealed significantly lower mtDNA copy number in both wb-PolgA^Mut/Mut^ and PolgA^Mut/Mut^ animals (Fig. 7D). Overall, these findings suggest similar levels of mitochondrial dysfunction in the novel wb-PolgA^Mut/Mut^ and orginal PolgA^Mut/Mut^ mouse models.

Next generation sequencing of the mtDNA of both models was also performed to assess mtDNA mutation load as well as specific trends in the mitochondrial genome. The results confirm similar elevated levels of mtDNA mutations in the wb-PolgA^Mut/Mut^ and PolgA^Mut/Mut^ mice (Fig. 7E). Both mutator models also have an increase in the transitions:transversions ratio as expected considering these are mainly due to replication error^36^ (Fig. 7F). Interestingly, the mutation pattern was also found to be nearly identical in both models, indicating that the nucleotide substitutions are being similarly made (Fig. 7G and H). Taken together, these data confirm that whole-body induction of the novel i-PolgA^Mut/Mut^ mouse recapitulates the original mtDNA mutator mouse model both genotypically and phenotypically.

Finally, to demonstrate spatial induction of the i-PolgA^Mut/Mut^ mouse, we generated a nervous system-specific PolgA^Mut/Mut^ mouse (ns-PolgA^Mut/Mut^). Using Cre-recombinase under control of a Nestin promoter, which is expressed in the central and peripheral nervous systems, we directed induction of the mutated polymerase-𝛾 in brain, but not in liver or ear biopsy (Fig. 7J). Future work will focus on validating and characterizing this nervous system-specific model to address questions on the impact of mtDNA mutations and resulting mitochondrial dysfunction on brain health and age-related cognitive decline.

In conclusion, this novel inducible mutator mouse expands the toolkit for studying the role of mitochondrial DNA mutations in aging and age-related diseases, where mitochondrial dysfunction has been implicated, including multiple forms of cancer, metabolic disorders, chronic inflammation, and neurodegenerative diseases. The spatial control of mtDNA mutations in our model enhances the ability to investigate their impact in specific tissues and cell types without affecting other tissues, allowing for a more precise dissection of mitochondrial dysfunction in organs such as the liver, muscle, heart^37–40^, and brain in both health and disease including neurodegeneration^22,30,41^. Studying the accumulation of these mutations in a tissue-specific manner will provide deeper insights into tissue-specific regulation of mtDNA mutation onset and clearance, and mitochondrial network dynamics. Furthermore, the temporal control offered by this model adds another layer of specificity, enabling researchers to assess how mitochondrial dysfunction affects health when mutations are induced at distinct time-points. This capability will be crucial for understanding the progression of mitochondrial dysfunction and its role in disease development over time.

## Materials and Methods

### Animals

The original mtDNA mutator strain, B6-Polg*^tm1.1(D257A)Lrsn^*, was obtained and homozygous mutator (PolgA^Mut/Mut^) mice and wild-type controls (PolgA^WT/WT^) were generated by intercrossing mice heterozygous for the mtDNA mutator PolgA^Mut^ allele and genotyped as previously described^19^. The novel inducible mutator mouse model, B6-Polg*^tm1.1(mPolgA)Ross^*, was generated in this study (genOway, Lyon, France) and homozygous mutator (i-PolgA^Mut/Mut^) animals were obtained by intercrossing mice heterozygous for the i-PolgA^Mut^ allele and genotyped accordingly. Induced whole-body mice, B6-Polg*^tm1.2(D257A)Ross^*, were generated using proprietary methods (genOway, Lyon, France) and homozygous whole-body mutator (wb-PolgA^Mut/Mut^) animals were obtained by intercrossing mice heterozygous for the wb-PolgA^Mut^ allele and genotyped accordingly. All mice were housed in the vivarium at the University of Rhode Island for at least two weeks prior to any handling. Mice were housed by sex with up to 5 mice per ventilated cage that contained tissues for nesting and access to a small hut. Animals were kept on a 12:12 light:dark cycle at 22 °C ± 1°C and 30-70% humidity. All mice received a standard diet (Teklad Global Soy Protein-Free (Irradiated) type 290X, Envigo, Indianapolis, IN, USA) and water ad libitum. Adequate measures were taken to minimize animal pain and discomfort. Investigation has been conducted in accordance with ethical standards and according to the Declaration of Helsinki and national and international guidelines and has been approved by the authors’ institutional review board (approval number AN1920-020).

### Generation of the novel inducible mtDNA mutator mouse

The novel inducible mtDNA mutator mouse model, B6-Polg*^tm1.1(mPolgA)Ross^* (i-PolgA^Mut/Mut^), was generated (genOway, Lyon, France) enabling inducible expression of the PolgA D257A mutant. The model based on the knock-in insertion, within mouse PolgA intron 3 of a transgene containing endogenous exon 3, followed by a cDNA encoding for wild-type PolgA, and containing endogenous 3’UTR and hGH polyA. Downstream of the hGH polyA a RoxP-flanked Neomycin resistance cassette was inserted, followed by mutant exon 3 harboring the D257A point mutation. The cDNA encodes for an optimized PolgA sequence in which cryptic splicing sites have been mutated in order to limit the risk of aberrant splicing events. This transgene was flanked by loxP sites, located within intron 3 and immediately downstream of the Neomycin cassette. Prior to Cre-recombinase action, the wild-type PolgA is expressed from the Knocked-in cDNA. Upon Cre-mediated recombination, the cDNA will be removed and mutant PolgA will be expressed. The homology arms were isolated by cloning from C57BL/6N mouse genomic DNA, which is isogenic to the ES cell line used for homologous recombination. The integrity of the targeting vector was assessed by full sequencing. Linearized targeting vector was transfected into C57BL/6J ES cells according to genOway’s standard electroporation procedures. G-418 resistant ES cell clones were subsequently validated by PCR, using primers hybridizing within and outside the targeted locus, and the whole recombined locus was sequenced to confirm the absence of genetic alteration. Recombined ES cell clones were microinjected into mouse blastocysts, which gave rise to male chimeras with a significant ES cell contribution. Breeding was established with C57BL/6N mice and produced the inducible PolgA mutant knock-in model devoid of the neomycin resistance cassette. Heterozygous mice were genotyped by PCR and further validated by full sequencing of the targeted locus, including the homology arms, and a few kbs upstream and downstream of the homology arms.

### Behavioral Experiments

Mice were acclimated to the testing room, a quiet, neutral environment kept at 22 °C ± 1°C, 30- 70% humidity, and ∼100 lux, in their home cage for 1 h prior to testing. All behavioral testing took place between 9:00 and 17:00 (light phase) by the same researcher, and the testing of mice from each experimental group and sex was randomized. The mice were transported to and from the apparatus in a non-transparent holding container that was cleaned with 70% ethanol after each use. Tests were conducted in the order described below.

### Open-Field Test (OF)

To assess exploratory behavior and spontaneous locomotion, the open-field behavioral assay was run using a multi-cage infrared-sensitive motion detection system (Fusion v6.5 SuperFlex, Omnitech electronics, Columbus, OH, USA). The mice were placed in darkened transparent chambers (40 x 40 x 30 cm) with a grid of infrared beams at the floor level and 7.5 cm above the floor. They were allowed to explore for 90 minutes while their movements were monitored in x-, y-, and z- planes and all horizontal and vertical activity recorded and analyzed using the v6.5 software system. The chambers were cleaned with 70% ethanol after each animal test period.

### Light-Dark Preference Test (LD)

To assess exploratory and anxiety-related behaviors animals were placed in the transparent locomotor chamber (40 x 40 x 30 cm) with a grid of infrared beams at floor level and 7.5 cm above the floor with an insert to divide the chamber into light and dark zones. The mice were allowed to explore for 30 minutes while their movements, as well as time in each zone, were monitored and recorded in x-, y-, and z- directions using the infrared system (Fusion v6.5 SuperFlex, Omnitech electronics, Columbus, OH, USA). The chambers and inserts were cleaned with 70% ethanol after each animal test period.

### Startle Reflex (SR)

To understand startle responses in the animals, the SR-LAB startle response system (SD Instruments, San Diego, CA, USA) was used. The system includes test cabinets with transparent animal enclosures (3.2 cm diameter) inside. Each test began with an acclimatization period of 5 min inside of the animal enclosure with exposure to white noise (70dB). There were 10 sessions with 11 trials each and within each session the animals were exposed to a range of auditory stimuli (20-120 dB) in a random order. Each stimuli lasts approximately 40 ms and the response from each animal were recorded for 150 ms. There was a mean of 15 s between each stimulus. The enclosures were cleaned with 70% ethanol between each animal test period.

### Pre-Pulse Inhibition (PPI)

To understand sensorimotor gating, the SR-LAB startle response system (SD Instruments, San Diego, CA, USA) was used for PPI. Mice were placed into the transparent animal enclosures (3.2 cm diameter) within the test cabinets and an acclimatization period with white noise (70dB) began the testing session. Each session began with 6 exposures to the startle alone, then 10 blocks with 6 different types of trials. These trials included a null trial where no stimuli was presented, startle trial of 120 dB, startle plus pre-plus with variable dB and pre-pulse alone. The trials were randomly interspersed, and each one started with a 50 ms null period to monitor movements at baseline followed by exposure to the pre-pulse alone and that response was recorded. Following this the startle stimuli were presented, and the responses recorded. The enclosures were cleaned with 70% ethanol between each animal test period. Percentage of Pre-Pulse Inhibition (PPI) was calculated using the following equation: % PPI = (1 - Startle response with pre-pulse/ Startle response without pre-pulse) x 100.

### Frailty Index

A 31-item frailty index was used as a non-invasive method to assess physical frailty of the mice^42^ as done previously^43^. Mice were presented at random to a blinded researcher who scored the animals on each of the index criteria. Measures included assessment of integument, the musculoskeletal, vestibulocochlear/auditory, ocular/nasal, digestive/urogenital, and respiratory systems, as well as overall discomfort of the mice. Based on this, animals were given a frailty index score as a measure of visible aging.

### Lifespan

Natural lifespan of wb-PolgA^WT/WT^, i-PolgA^Mut/Mut^, and wb-PolgA^Mut/Mut^ mice were estimated by a survival study as previously described^44^. Mice were euthanized upon any sign of disease or distress. Animals were monitored for symptoms such as hunched shoulders, inactivity, failure to eat or drink, unresponsive to external stimuli, loss of more than 20% of their body weight, lack of grooming behavior, etc.

### Tissue Preparation

Mice were anesthetized via intraperitoneal injection of sodium pentobarbital (200 mg/kg) and cervically dislocated. Blood was collected via cardiac puncture and brain, heart, gastrocnemius muscle, kidney, liver, and spleen tissues were quickly frozen on dry ice or post-fixed in 10% formalin (Epredia, Portsmouth, NH, USA) for 24 h at 4°C followed by 30% sucrose in 1x PBS. Tissue was then embedded as necessary (Tissue-Plus OCT compound, Fisher Scientific, Waltham, MA, USA) and stored at -80°C until use.

### Western Blot

Liver, spleen, and cerebellum samples from male i-PolgA^WT/WT^ and i-PolgA^Mut/Mut^ mice were lysed in RIPA buffer (50 mM Tris-HCl pH 7.4, 150 mM NaCl, 0.5% deoxycholic acid, 0.1% sodium dodecyl sulfate, 2 mM EDTA, 1% Triton X-100) containing a proteinase (1:100, 78,438, Halt, Thermo Scientific, Fremont, CA, USA) and phosphatase (1:100, P0044, Sigma Aldrich) inhibitor cocktail. The samples were then incubated on ice for 30 min and sonicated (QSonica, Newtown, CT, USA) for 3 min (30–30 pulse) at 4 °C with 30% amplitude and centrifuged at 10,000 *g* for 10 min. To determine protein concentration, a BCA protein assay kit (23225 Thermo Scientific, Fremont, CA, USA) was used according to the manufacturer’s instructions. The samples were then prepared by mixing the lysate with loading buffer (1610747, Bio-Rad, Hercules, CA, USA) plus 100 mM DTT and incubated at 98 °C for 10 min. Protein mixture was then separated in an SDS-PAGE precast 4-20% gradient gel (5671094, Bio-Rad, Hercules, CA, USA). Samples were then blotted on a 0.45 µm nitrocellulose membrane (10600002, Cytiva, Marlborough, MA, USA) using transfer buffer (25 mM Tris-HCl pH 8.3, 190 mM glycine 20% methanol). The membrane was blocked (0.1% TBS-Tween with 5% skim milk) for 1 h and probed with rabbit anti-PolgA primary (1:1,000, MA5-37845, Invitrogen, Waltham, MA, USA) and goat anti-rabbit-HRP secondary antibodies in 0.1% TBS-Tween (1:3000, 1706515, Bio-Rad, Hercules, CA, USA). Immunocomplexes were detected by chemiluminescence (SuperSignal Chemiluminescence Substrate, Thermo Scientific, Fremont, CA, USA) and visualized (ChemiDoc, BioRad, Hercules, CA, USA). Images were processed and quantified using appropriate software (FIJI v2.1.0/1.53c, Madison, WI, USA).

### Quantitative Real-Time PCR (qPCR) for PolgA Expression

Liver tissue from i-PolgA^WT/WT^ and i-PolgA^Mut/Mut^ were lysed and RNA was extracted and purified (ZYmo Direct-zol RNA MiniPrep Plus Kit R2070, Zymo Research, Irvine, CA, USA) according to the manufacturer’s protocol. A spectrophotometer (NanoDrop, ND-2000, Thermo Scientific, Fremont, CA, USA) was used to determine RNA concentration. To synthesize cDNA, reverse transcription was run (Lunascript^®^ RT Supermix Kit E3010, New England BioLabs, Inc. Ipswich, MA, USA) according to the manufacturer’s protocol. Following this, qPCR reactions were run using a SYBR green-based master mix (LUNA^®^ qPCR Mastermix M3003, New England BioLabs Inc.) based on the manufacturer’s instructions on an appropriate instrument (Lightcycler 96, Roche, Basel, Switzerland). The reaction conditions were 95 °C for 60 s, 95 °C for 15 s with an annealing temperature of 61 °C, and melt curve of 65–90 °C. Primer sequences were as follows *PolgA*_For_: CTG CCG CAG AAC GGG AAG C; *PolgA*_Rev_: CTT TGG GCT CCA GCT TGA C.

### Quantitative Real-Time PCR (qPCR) for Copy Number

Copy number was determined based on mitochondrial gene ND2, normalized to nuclear gene 18s. Samples of gastrocnemius muscle from male i-PolgA^WT/WT^, i-PolgA^Mut/Mut^, PolgA^Mut/Mut^, and wb- PolgA^Mut/Mut^ animals (N=6 per group) were lysed and DNA was extracted as described above using phenol-chloroform extraction^23^. To determine DNA concentration a spectrophotometer (NanoDrop, ND-2000, Thermo Scientific, Fremont, CA, USA) was used. qPCR reactions were run using TaqMan based master mix (TaqMan^TM^ Fast Advanced Master Mix, 4444556, Thermo Fisher Scientific, Waltham MA, USA) according to manufacturer’s instructions with an appropriate instrument (Lightcycler 96, Roche, Basel, Switzerland). Primers for 18s (TaqMan Pre-Developed Assay Reagents, Euk 18srRNA, 4333760T, Thermo Fisher, Waltham MA, USA) and ND2 (TaqMan Gene Expression Assays, 4448484 PL, Thermo Fisher, Waltham MA, USA) were used with FAM and VIC dyes respectively. The reaction conditions were 95 °C for 20 s, 95 °C for 3 s with annealing temperature of 60 °C for 30 s × 40.

### Complex I Activity Assay

To assess mitochondrial function, mitochondrial respiratory Complex I activity was assessed in liver tissue according to manufacturer’s instructions (ab109721, Abcam, Cambridge, MA, USA). Liver was homogenized in cold 1x PBS, BCA was performed to quantify protein as described above, and detergent was added to extract protein. Samples were then centrifuged at 16,000 x *g* for 20 min and supernatant was collected. Protein levels were normalized, diluted to appropriate range and loaded into wells with incubation solution and allowed to incubate for 3 h at RT, shaking at 400 rpm. The plate was then washed, an assay solution was added, and samples were read using an appropriate instrument (BioTek Synergy HTX Reader, Agilent, Santa Clara, CA, USA) at 450 nm, shaking and recording every minute for 40 min.

### Transmission Electron Microscopy

Tissue fixation was done via transcardial perfusion of animals with 10% formalin. Gastrocnemius muscle was collected and transversely cut into small pieces, stored in Karnvosky fixative: 2.5% glutaraldehyde (16019, Electron Microscopy Sciences, Hatfield, PA, USA) in 0.15 M Na Cacodylate (11650, Electron Microscopy Sciences, Hatfield, PA, USA), and 2% paraformaldehyde (15710, Electron Microscopy Sciences, Hatfield, PA, USA) plus 2 mM calcium chloride (C79- 500, Fisher Scientific, Waltham, MA, USA) at 4 °C. Samples were then washed, processed, embedded^45^ and imaged with assistance from The Leduc Imaging Facility (Brown University, Providence, RI, USA).

### Cytochrome C Oxidase/Succinate Dehydrogenase stain (COX/SDH)

To visualize mitochondrial function, fresh-frozen embedded liver tissue from PolgA^Mut/Mut^, wb- PolgA^Mut/Mut^, i-PolgA^Mut/Mut^, and i-PolgA^WT/WT^ were cryosectioned (Leica BioSystems, CM1950, Wetzlar, Germany) at 14 µm at -21°C and mounted directly onto slides (VWR Colorfrost Plus, Radnor PA, USA). Slides were then incubated with COX staining solution for 20 min at room temperature as done previously^46^.

### Microscopy

Brightfield microscopy was used to evaluate the enzyme histochemistry results (Leica DMi8 microscope, Wetzlar, Germany). Images were processed using appropriate software (FIJI v2.1.0/1.53c, Madison, WI, USA).

### Next Generation Sequencing (NGS)

Samples of liver tissue were lysed overnight at 55 °C shaking gently (400-600 rpm) in digestion buffer containing Tris-HCl (10mM), EDTA (25mM), SDS (0.5%), NaCl (100mM), and proteinase K (200 µg/ml, Sigma Aldrich, Burlington, MA, USA). The samples were then vortexed and centrifuged (8,000 *g* for 15 min), then 0.6 ml of supernatant was mixed with 0.6 ml of phenol/chloroform/isoamyl alcohol (25:4:1 PCIAA, P2069, Sigma Aldrich, Burlington, MA, USA). Following vigorous mixing, the samples were centrifuged (8,000 *g* for 15 min), and 0.45-0.5 ml of supernatant was removed and combined with an equal volume of chloroform (C2432, Sigma Aldrich, Burlington, MA, USA). After mixing, the samples were centrifuged, and 0.4 ml of the supernatant was added to 0.04 ml NaAc (3M) and 0.44 ml isopropanol. The samples were then kept at -20 °C for 10-20 min, centrifuged (8000 *g* 15 min) and supernatant was discarded taking care to not disturb the DNA pellet. The pellet was washed was 1 ml of 70% ethanol, air dried and then resuspended in 0.4 ml tris-EDTA buffer. Spectrophotometer (Nanodrop, Thermo Scientific, Fremont, CA, USA) and PCR were used to confirm presence and quality of DNA. Amplification of the mitochondrial genome was done using three primers sets to amplify three ∼6 kb regions covering the entire mitochondrial genome^47^ using Phusion High-Fidelity DNA polymerase (F530L, Thermo Scientific, Fremont, CA, USA) and amplified with an appropriate instrument (T100 Thermal Cycler, BioRad, Hercules, CA, USA). The reactions conditions were were 98 °C for 30 s, 98 °C for 10 s with optimal annealing temperature varying by primer for 30 s × 30. Primer sequences were as follows *mtDNA1_For_:* CCA TAA ACA CAA AGG TTT GGT CC; *mtDNA1*_Rev_: GGT TGA CCT AAT TCT GCT CGA AT, *mtDNA2_For_:* GGA AAC TGA CTT GTC CCA; *mtDNA2_Rev_*: GCG TAA GCA GAT TGA GCT, *mtDNA3_For_:* CGC CTA CTC CTC AGT TAG C; *mtDNA3_Rev_*: AGA GTT TTG CAC GGA. Library preparation was done using the KAPA HyperPlus Kit (Roche, Indianapolis, IN, USA) according to the manufacturer’s instructions. The DNA library was then checked for quality and sequenced using the MiSeq Illumina using the MiSeq v3 Reagent Kit (Illumina, San Diego, CA, USA). Sequence data were analyzed using a Snakemake implementation of a GATK human mitochondrial variant analysis pipeline (*gatk.broadinstitute.org/hc/en-us/articles/4403870837275Mitochondrial-short-variant-discovery-SNVsIndels-)* generalized to use any organism. In summary, the pipeline aligns pre- processed reads to the mm39 genome, subsets mitochondrial reads to account for nuclear mitrochondrial DNA (NUMT’s), realigns mitochondrial reads to two versions of the mm39 mitochondrial genome (one coordinate shifted 180°) to ensure coverage of the mitochondrial control region, deduplicates the reads, calls variants, merges the shifted and unshifted vcf files together, and finally filters the variants.

### Statistical Analysis

Data are presented as mean values (M) with SEM. Statistical analyses, unpaired t-tests, one-way ANOVAs, or two-way ANOVAs with Tukey post-hoc multiple comparisons were performed using appropriate software (GraphPad Prism v.10, San Diego, CA, USA). Significances are denoted as \**p* < .05, \*\**p* < .01, \*\*\**p* < .001, and \*\*\*\**p* < .0001, with α *p* < .10 as trending.

## Author Contributions

J.M.R., G.C., and L.O. conceived the idea to generate the novel inducible mtDNA mutator mouse model (i-PolgA^Mut/Mut/^). J.M.R. provided funding to generate the novel inducible mtDNA mutator (i-PolgA^Mut/Mut/^) and whole-body induced mtDNA mutator (wb-PolgA^Mut/Mut^) models. E.S., V.D., C.B. at genOway generated the i-PolgA^Mut/Mut^ and the induced wb-PolgA^Mut/Mut^ mice. H.T.W., G.C. and J.M.R. designed the experiments. H.T.W., S.B, L.G., and B.G. performed the experiments, and H.T.W., G.C., and C.H. analyzed the data. H.T.W., G.C. and J.M.R. wrote the manuscript and all authors have read and agreed to the published version of the manuscript.

## Acknowledgements

The novel inducible mtDNA mutator (i-PolgA^Mut/Mut/^) and whole-body induced mtDNA mutator (wb-PolgA^Mut/Mut^) models were generated using a monetary gift given to J.M.R. from Konung Gustaf V:s och Drottning Victorias Frimurarestiftelse. This research was supported by the National Institutes of Health Office of the Director under grant number R21OD037651 (to J.M.R., G.C.), the College of Pharmacy Seed Grant at the University of Rhode Island (to G.C.), the First-Year Doctoral Fellowship at the University of Rhode Island (to H.T.W.), and the George and Anne Ryan Institute for Neuroscience at the University of Rhode Island (to G.C. and J.M.R.). Research was facilitated by the use of equipment and services available through the Rhode Island Institutional Development Award (IDeA) Network of Biomedical Research Excellence from the National Institute of General Medical Sciences of the National Institutes of Health under grant number P20GM103430 through the Centralized Research Core facility and the Molecular Informatics Core (RRID:SCR_017685). We thank NH INBRE for bioinformatic consultation, and Geoff Williams and the Leduc Bioimaging Facility at Brown University for their technical support.

## Institutional Review Board Statement

The animal study protocol (#AN1920-020) was approved by the Institutional Review Board of University of Rhode Island, under J.M.R. and G.C.

## Data Availability Statement

Data generated from this study are available upon request.

