## Supplementary Figures for "A novel inducible mtDNA mutator mouse model to study mitochondrial dysfunction with temporal and spatial control"

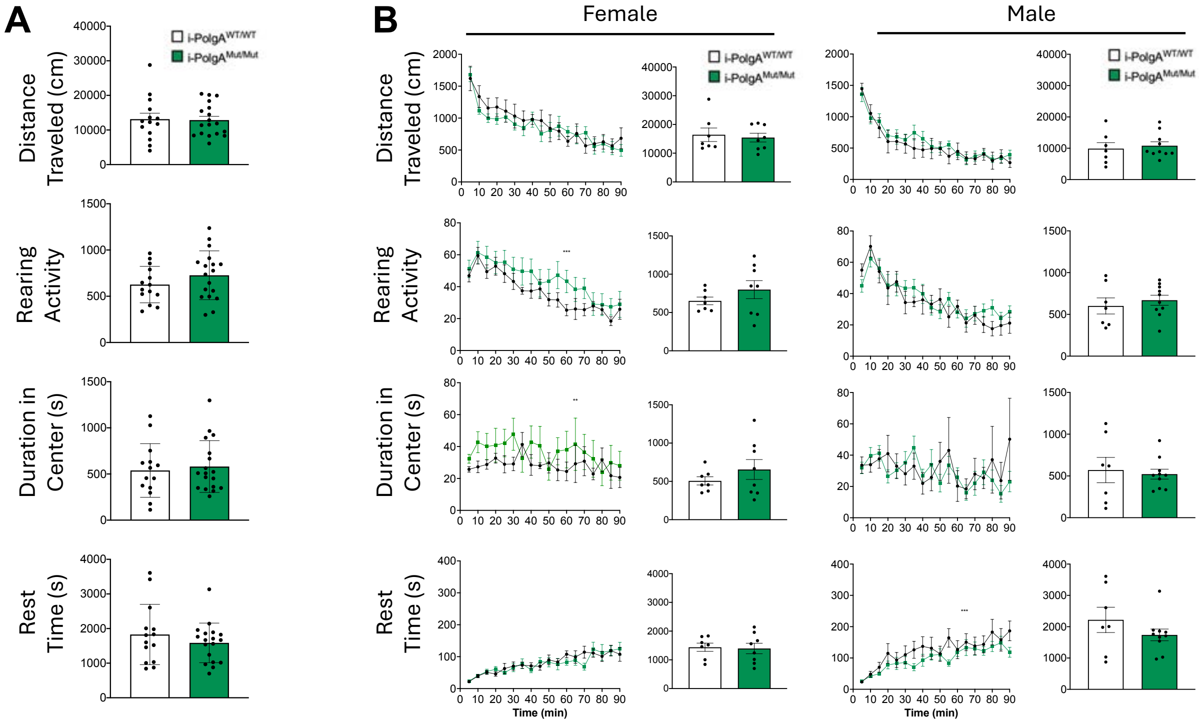


**Supplementary Figure S1. Open-field behavioral assay in 9 months old i-PolgA mice. (A)** Total spontaneous locomotion and exploratory activity over 90 minutes in i-PolgA$^{WT/WT}$ (N=14) and i-PolgA$^{Mut/Mut}$ (N=18) mice. **(B)** Spontaneous locomotion and exploratory activity in female i-PolgA$^{WT/WT}$ (N=7) and i-PolgA$^{Mut/Mut}$ (N=8) and male i-PolgA$^{WT/WT}$ (N=7) and i-PolgA$^{Mut/Mut}$ (N=10) animals. Significances were determined with two-way ANOVA with post-hoc analysis and ** $p < 0.01$, *** $p < 0.001$.

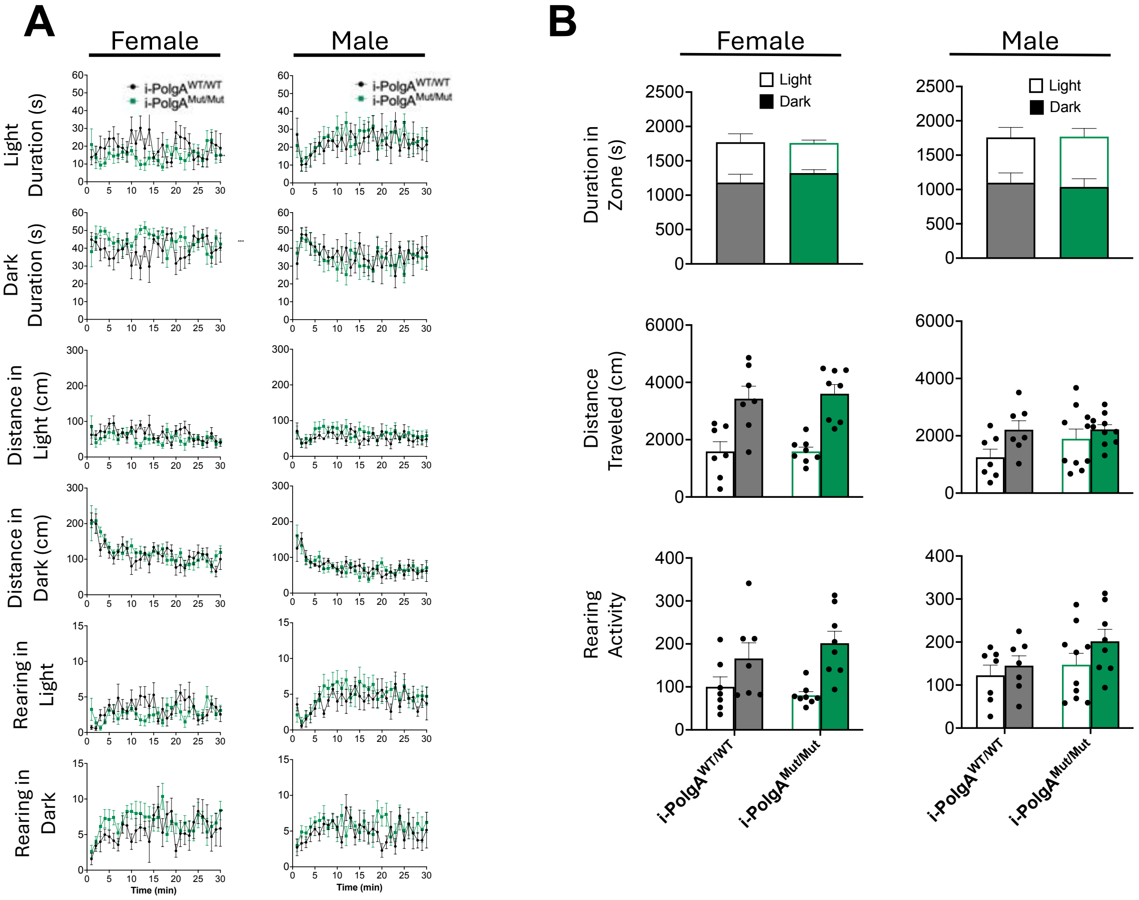


**Supplementary Figure S2. Light-dark behavioral assay in 9 months old i-PolgA mice. (A)** Exploratory behavior and anxiety-like behavior in the light-dark preference assay in female i-PolgA$^{WT/WT}$ (N=7) and i-PolgA$^{Mut/Mut}$ (N=8) and male i-PolgA$^{WT/WT}$ (N=7) and i-PolgA$^{Mut/Mut}$ (N=10) animals. **(B)** Total exploratory behavior and anxiety-like behavior over 30 minutes in the light-dark preference assay in female i-PolgA$^{WT/WT}$ (N=7) and i-PolgA$^{Mut/Mut}$ (N=8) and male i-PolgA$^{WT/WT}$ (N=7) and i-PolgA$^{Mut/Mut}$ (N=10) animals. Significances were determined with two-way ANOVA with post-hoc analysis and *** $p < 0.001$.

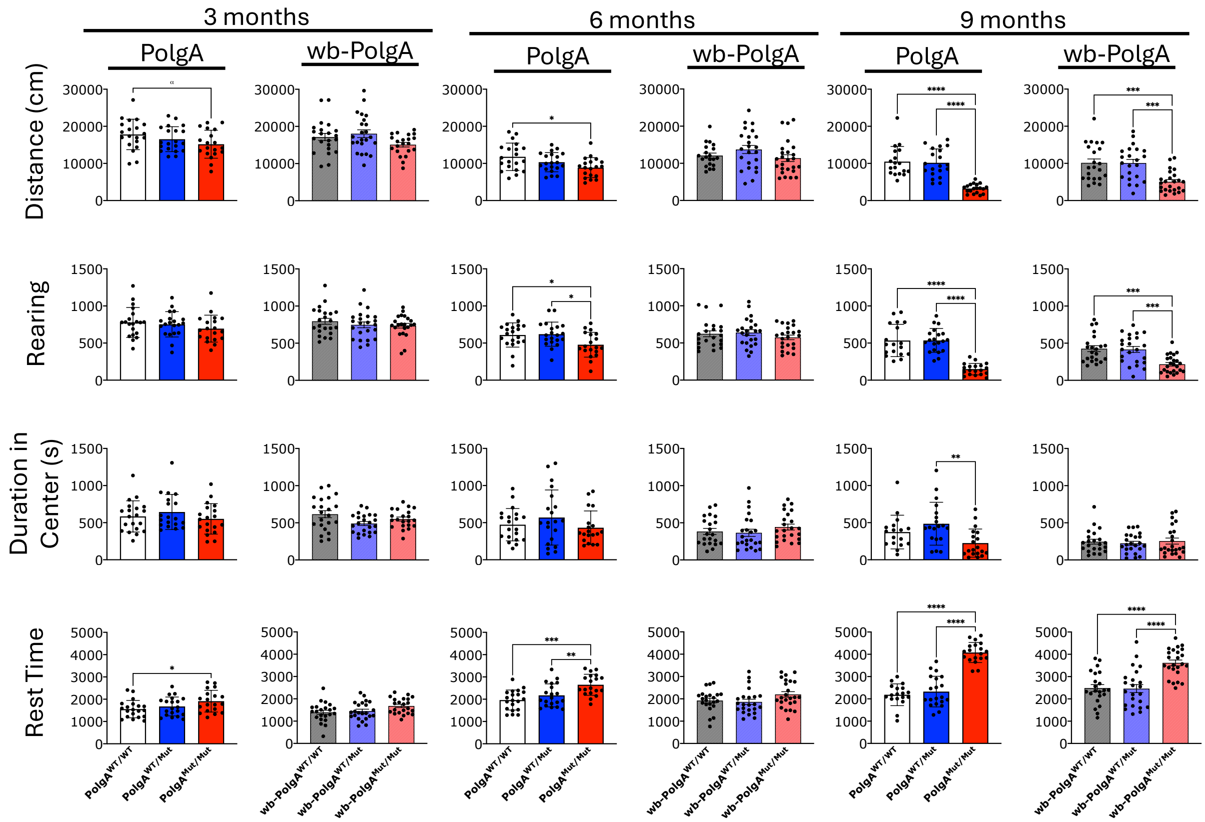


**Supplementary Figure S3. Open-field behavioral assay in 3, 6, and 9 month-old PolgA and wb-PolgA mice.** Total spontaneous locomotion and exploratory activity over 90 minutes in PolgA$^{WT/WT}$ (N=20), PolgA$^{WT/Mut}$ (N=20), PolgA$^{Mut/Mut}$ (N=20), and in wb-PolgA$^{WT/WT}$ (N=22), wb-PolgA$^{WT/Mut}$ (N=23), wb-PolgA$^{Mut/Mut}$ (N=24) animals. Both PolgA$^{Mut/Mut}$ and wb-PolgA$^{Mut/Mut}$ animals exhibit altered behavior including increased rest and decreased rearing and distance traveled. Significances were determined by one-way ANOVA with post-hoc analysis; * $p < 0.05$, ** $p < 0.01$, *** $p < 0.001$, **** $p < 0.0001$, and α $p < 0.10$ as trending.

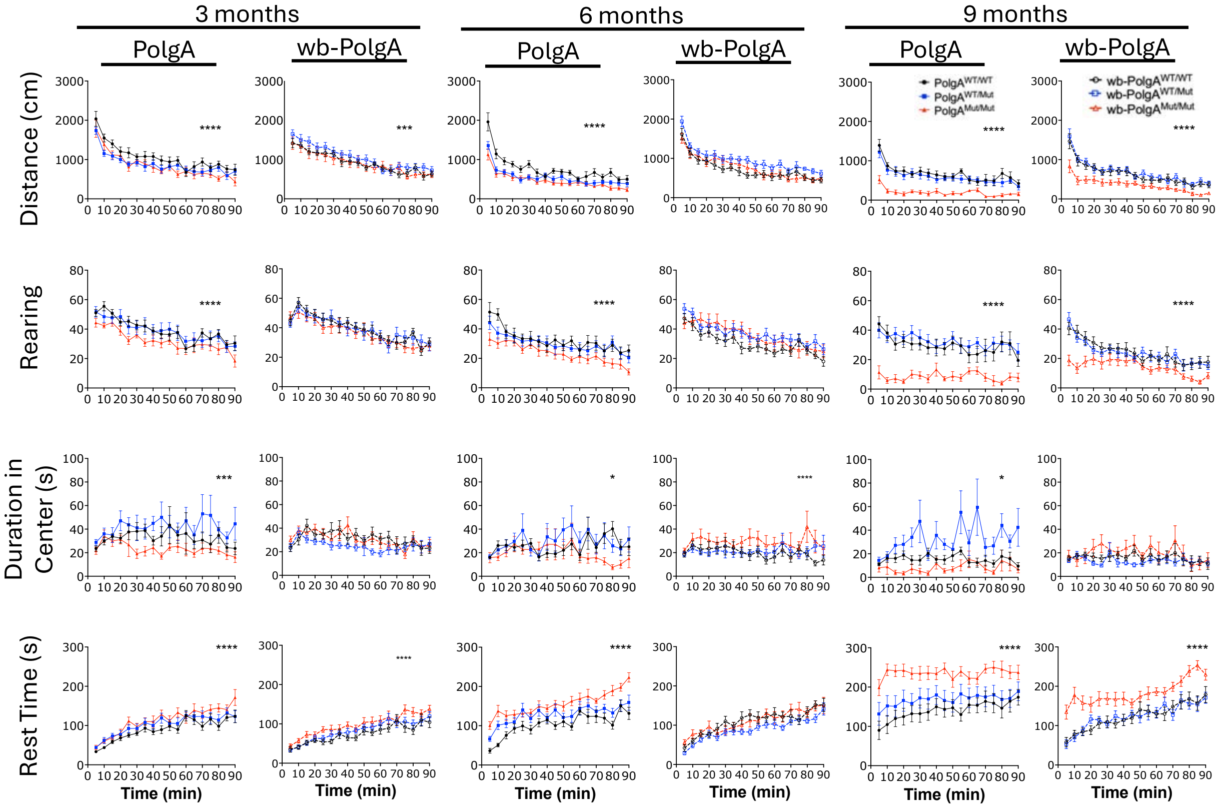


**Supplementary Figure S4**. **Open-field behavioral assay in 3, 6, and 9 months old female PolgA and wb-PolgA mice.** Spontaneous locomotion and exploratory activity in female PolgA$^{WT/WT}$ (N=10), PolgA$^{WT/Mut}$ (N=10), PolgA$^{Mut/Mut}$ (N=10), and in wb-PolgA$^{WT/WT}$ (N=11), wb-PolgA$^{WT/Mut}$ (N=12), wb-PolgA$^{Mut/Mut}$ (N=12) animals. Both PolgA$^{Mut/Mut}$ and wb-PolgA$^{Mut/Mut}$ animals exhibit altered behavior including increased rest and decreased rearing and distance traveled. Significances were determined by two-way ANOVA with post-hoc analysis and * $p < 0.05$, ** $p < 0.01$, *** $p < 0.001$, **** $p < 0.0001$.

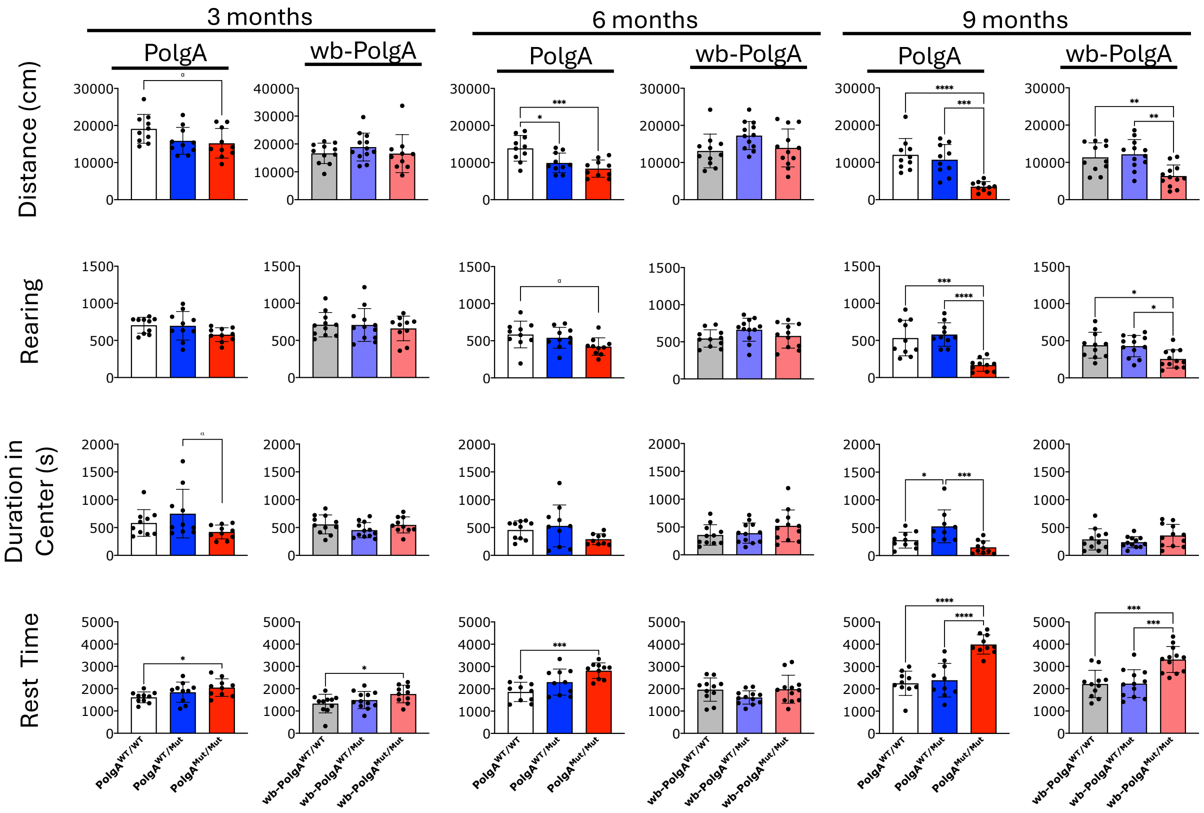


**Supplementary Figure S5**. **Open-field behavioral assay in 3, 6, and 9 months old female PolgA and wb-PolgA mice.** Total spontaneous locomotion and exploratory activity over 90 minutes in female PolgA$^{WT/WT}$ (N=10), PolgA$^{WT/Mut}$ (N=10), PolgA$^{Mut/Mut}$ (N=10), and in wb-PolgA$^{WT/WT}$ (N=11), wb-PolgA$^{WT/Mut}$ (N=12), wb-PolgA$^{Mut/Mut}$ (N=12) animals. Both PolgA$^{Mut/Mut}$ and wb-PolgA$^{Mut/Mut}$ animals exhibit altered behavior including increased total rest and decreased total rearing and total distance traveled. Significances were determined by one-way ANOVA with post hoc analysis; * $p < 0.05$, ** $p < 0.01$, *** $p < 0.001$, **** $p < 0.0001$, and $\alpha$ $p < 0.10$ as trending.

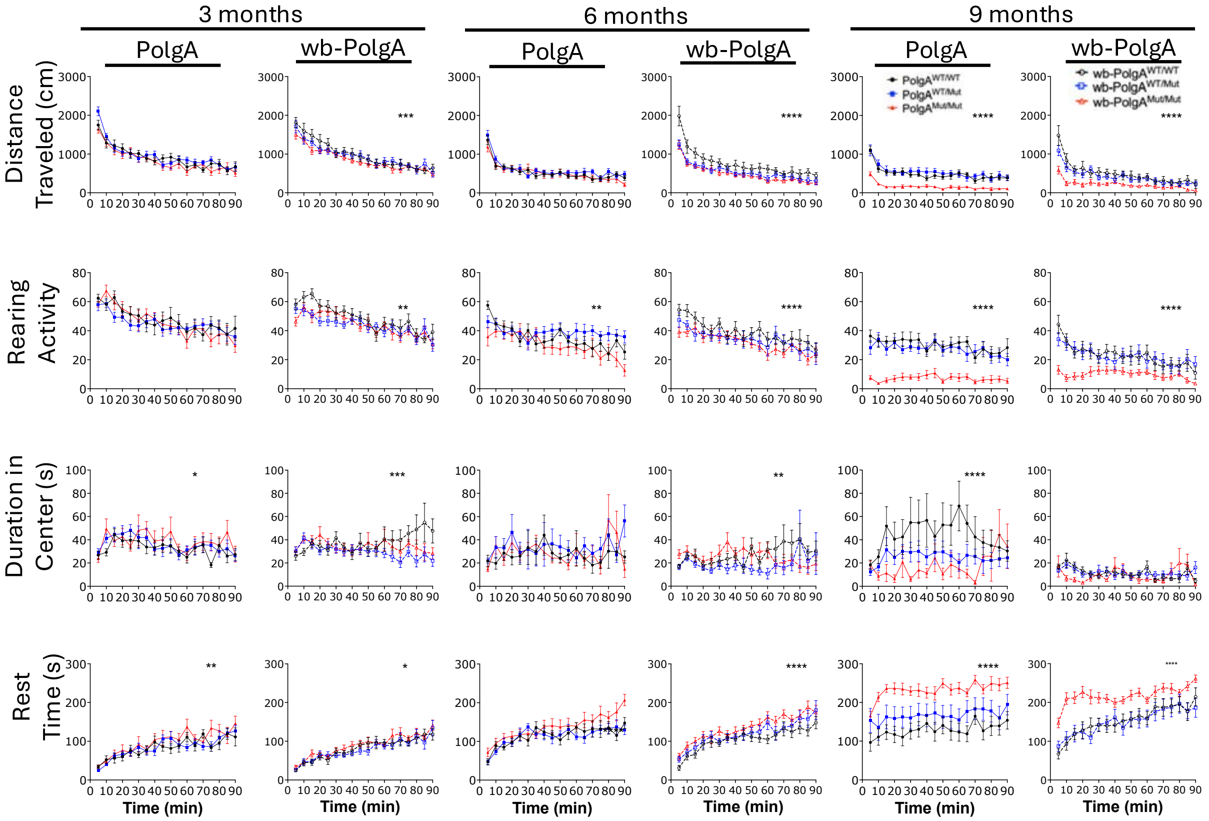


**Supplementary Figure S6**. **Open-field behavioral assay in 3, 6, and 9 months old male PolgA and wb-PolgA mice.** Spontaneous locomotion and exploratory activity in male PolgA$^{WT/WT}$ (N=10), wb-PolgA$^{WT/Mut}$ (N=10), PolgA$^{Mut/Mut}$ (N=10), and in wb-PolgA$^{WT/WT}$ (N=11), wb-PolgA$^{WT/Mut}$ (N=11), wb-PolgA$^{Mut/Mut}$ (N=12) animals. Both PolgA$^{Mut/Mut}$ and wb-PolgA$^{Mut/Mut}$ animals exhibit altered behavior including increased rest and decreased rearing and distance traveled. Significances were determined by two-way ANOVA with post-hoc analysis and * $p < 0.05$, ** $p < 0.01$, *** $p < 0.001$, **** $p < 0.0001$.

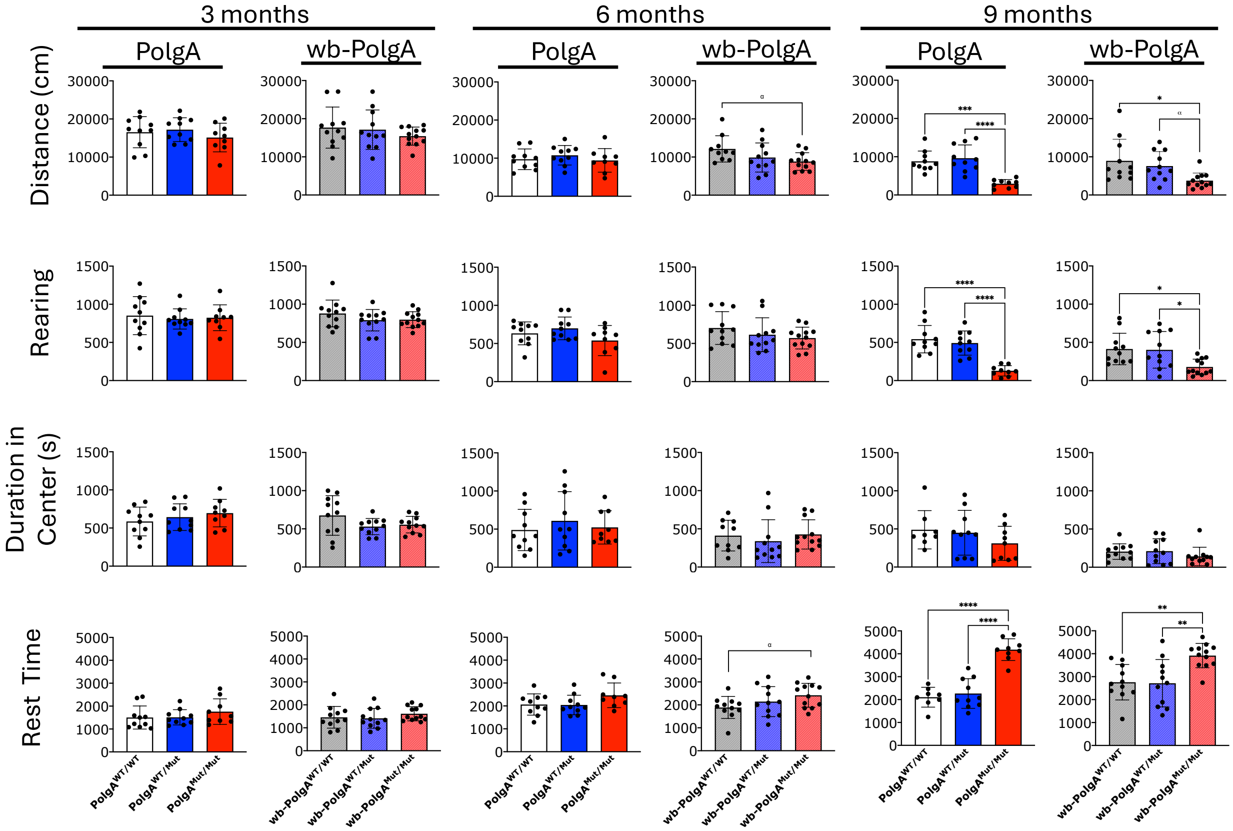


**Supplementary Figure S7**. **Open-field behavioral assay in 3, 6, and 9 months old male PolgA and wb-PolgA mice.** Total spontaneous locomotion and exploratory activity in male PolgA$^{WT/WT}$ (N=10), PolgA$^{WT/Mut}$ (N=10), PolgA$^{Mut/Mut}$ (N=10), and in wb-PolgA$^{WT/WT}$ (N=11), wb-PolgA$^{WT/Mut}$ (N=11), wb-PolgA$^{Mut/Mut}$ (N=12) animals. Both PolgA$^{Mut/Mut}$ and wb-PolgA$^{Mut/Mut}$ animals exhibit altered behavior including increased total rest and decreased total rearing and total distance traveled. Significances were determined by one-way ANOVA with post-hoc analysis; * $p < 0.05$, ** $p < 0.01$, *** $p < 0.001$, **** $p < 0.0001$, and α $p < 0.10$ as trending.

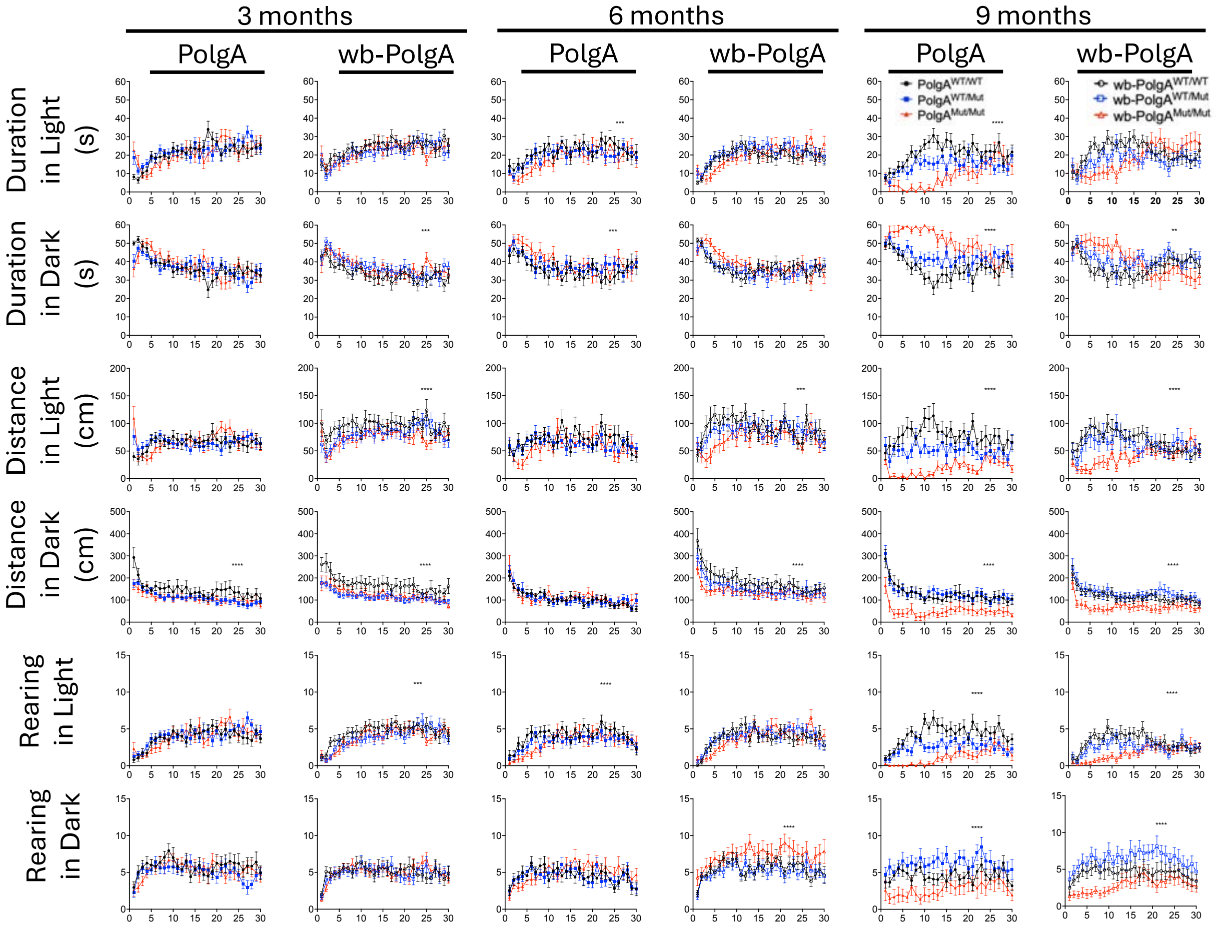


**Supplementary Figure S8. Light-dark behavioral assay in 3, 6, and 9 months old in PolgA and wb-PolgA mice.** Exploratory behavior and anxiety-like behavior in the light-dark preference assay in PolgA$^{WT/WT}$ (N=20), PolgA$^{WT/Mut}$ (N=20), PolgA$^{Mut/Mut}$ (N=20), and in wb-PolgA$^{WT/WT}$ (N=22), wb-PolgA$^{WT/Mut}$ (N=23), wb-PolgA$^{Mut/Mut}$ (N=24) animals. Both PolgA$^{Mut/Mut}$ and wb-PolgA$^{Mut/Mut}$ animals exhibit altered behavior including decreased rearing and distance traveled. Significances were determined by two-way ANOVA with post-hoc analysis; * $p < 0.05$, ** $p < 0.01$, *** $p < 0.001$, **** $p < 0.0001$, and α $p < 0.10$ as trending.

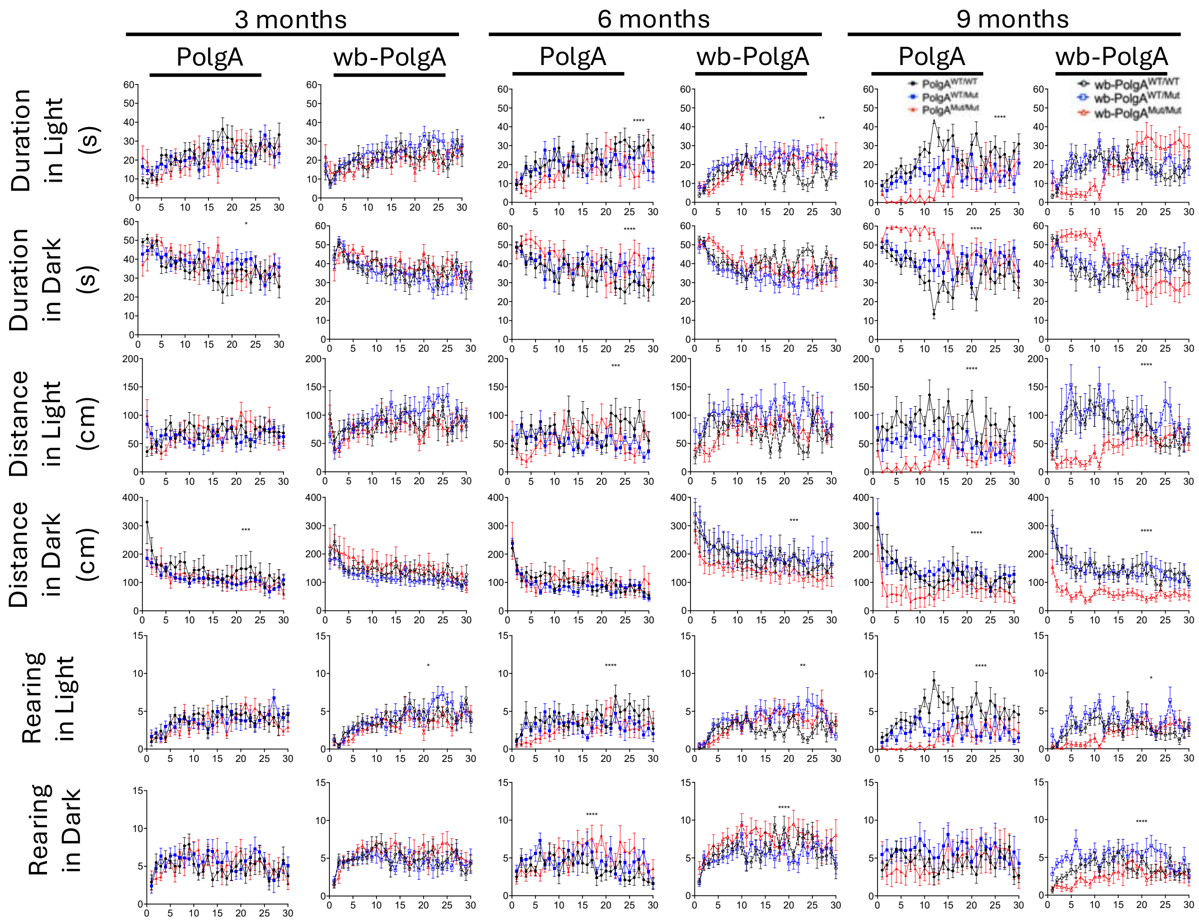


**Supplementary Figure S9. Light-dark behavioral assay in 3, 6, and 9 months old in female PolgA and wb-PolgA mice.** Exploratory behavior and anxiety-like behavior in the light-dark preference assay in female PolgA$^{WT/WT}$ (N=10), PolgA$^{WT/Mut}$ (N=10), PolgA$^{Mut/Mut}$ (N=10), and in wb-PolgA$^{WT/WT}$ (N=11), wb-PolgA$^{WT/Mut}$ (N=12), wb-PolgA$^{Mut/Mut}$ (N=12) animals. Both PolgA$^{Mut/Mut}$ and wb-PolgA$^{Mut/Mut}$ animals exhibit altered behavior including decreased rearing and distance traveled. Significances were determined by two-way ANOVA with post-hoc analysis and * $p < 0.05$, ** $p < 0.01$, *** $p < 0.001$, **** $p < 0.0001$.

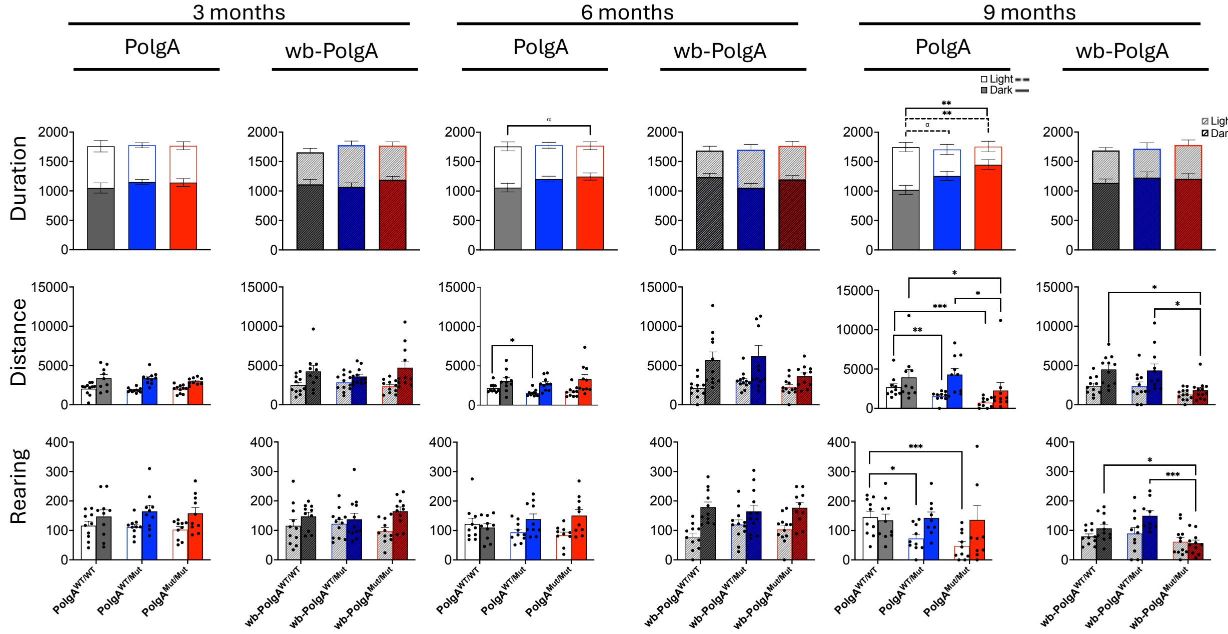


**Supplementary Figure S10. Light-dark behavioral assay in 3, 6, and 9 months old female PolgA and wb-PolgA mice.** Total exploratory behavior and anxiety-like behavior over 30 minutes in the light-dark preference assay in female PolgA$^{WT/WT}$ (N=10), wb-PolgA$^{WT/Mut}$ (N=10), PolgA$^{Mut/Mut}$ (N=10), and in wb-PolgA$^{WT/WT}$ (N=11), wb-PolgA$^{WT/Mut}$ (N=12), wb-PolgA$^{Mut/Mut}$ (N=12) animals. Both PolgA$^{Mut/Mut}$ and wb-PolgA$^{Mut/Mut}$ animals exhibit altered behavior including decreased rearing and distance traveled. Significances were determined by one-way ANOVA with post-hoc analysis; * $p < 0.05$, ** $p < 0.01$, *** $p < 0.001$, **** $p < 0.0001$; and α $p < 0.10$ as trending.

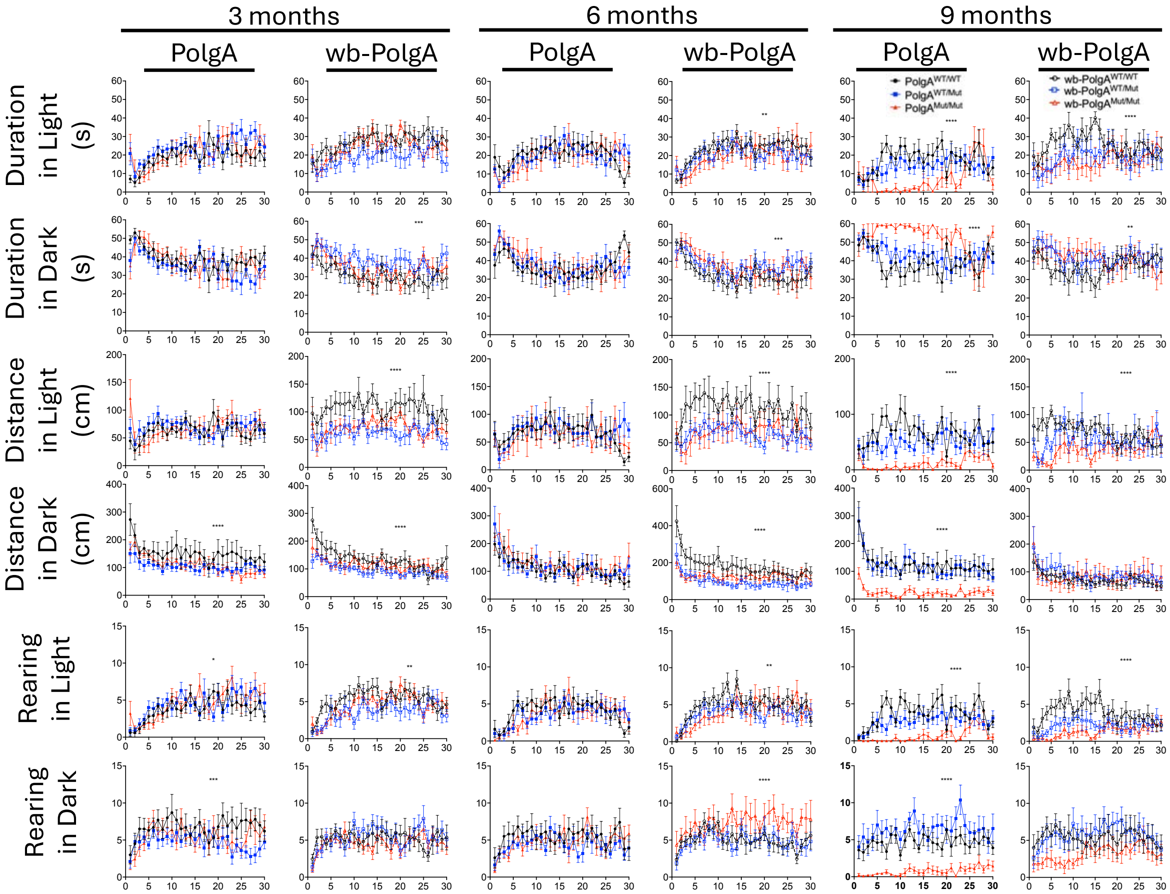


**Supplementary Figure S11. Light-dark behavioral assay in 3, 6, and 9 months old male PolgA and wb-PolgA mice.** Exploratory behavior and anxiety-like behavior in the light-dark preference assay in male PolgA$^{WT/WT}$ (N=10), PolgA$^{WT/Mut}$ (N=10), PolgA$^{Mut/Mut}$ (N=10), and in wb-PolgA$^{WT/WT}$ (N=11), wb-PolgA$^{WT/Mut}$ (N=11), wb-PolgA$^{Mut/Mut}$ (N=12) animals. Both PolgA$^{Mut/Mut}$ and wb-PolgA$^{Mut/Mut}$ animals exhibit altered behavior including decreased rearing and distance traveled. Significances were determined by two-way ANOVA with post-hoc analysis and * $p < 0.05$, ** $p < 0.01$, *** $p < 0.001$, **** $p < 0.0001$.

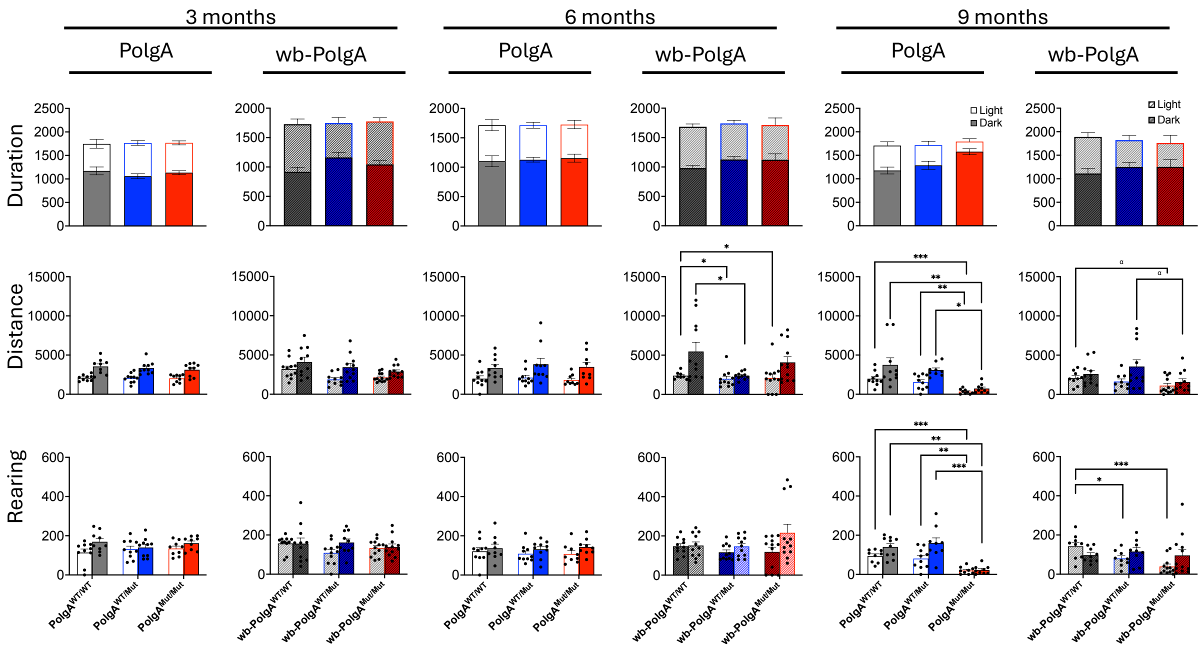


**Supplementary Figure S12. Light-dark behavioral assay in 3, 6, and 9 months old male PolgA and wb-PolgA mice.** Total exploratory behavior and anxiety-like behavior over 30 minutes in the light-dark preference assay in male PolgA$^{WT/WT}$ (N=10), PolgA$^{WT/Mut}$ (N=10), PolgA$^{Mut/Mut}$ (N=10), and in wb-PolgA$^{WT/WT}$ (N=11), wb-PolgA$^{WT/Mut}$ (N=11), wb-PolgA$^{Mut/Mut}$ (N=12) animals. Both PolgA$^{Mut/Mut}$ and wb-PolgA$^{Mut/Mut}$ animals exhibit altered behavior including decreased rearing and distance traveled. Significances were determined by one-way ANOVA with post-hoc analysis; * $p < 0.05$, ** $p < 0.01$, *** $p < 0.001$, **** $p < 0.0001$, and $\alpha$ $p < 0.10$ as trending.
